# Metabolomic Response to Non-Steroidal Anti-Inflammatory Drugs

**DOI:** 10.1101/2024.11.26.625478

**Authors:** Soumita Ghosh, Nick Lahens, Kayla Barekat, Soon-Yew Tang, Katherine N. Theken, Emanuela Ricciotti, Arjun Sengupta, Robin Joshi, Frederic D. Bushman, Aalim Weljie, Tilo Grosser, Garret A FitzGerald

## Abstract

Non-steroidal anti-inflammatory drugs (NSAIDs) are popular choices for the mitigation of pain and inflammation; however, they are accompanied by side effects in the gastrointestinal and cardiovascular systems. We compared the effects of naproxen, a traditional NSAID, and celecoxib, a cyclooxygenase -2 (Cox-2) inhibitor, in humans. Our findings showed a decrease in tryptophan and kynurenine levels in plasma of volunteers treated with naproxen. We further validated this result in mice. Additionally, we find that the depression of tryptophan was independent of both Cox-1 and Cox-2 inhibition, but rather was due to the displacement of bound tryptophan by naproxen. Supplementation of tryptophan in naproxen-treated mice rescued fecal blood loss and inflammatory gene expression driven by IL-1β in the heart.

## Introduction

Non-steroidal anti-inflammatory drugs (NSAIDs) are antipyretic^1^, analgesic^2^, and anti-inflammatory drugs, used for managing pain and inflammation; they are amongst the most consumed drugs worldwide. They act by inhibiting the cyclooxygenase (Cox) enzymes that metabolize arachidonic acid (AA) to prostaglandins (PGs) and thromboxane (Tx)^3^. These prostanoids are mediators of inflammation and fever, playing a role in the pathophysiology of many diseases^4^. Although NSAIDs offer a non-addictive option in the management of pain, NSAIDs cause gastrointestinal and cardiovascular adverse effects^5,6,7^.

Here, we compared the effects of two NSAIDs, celecoxib, an inhibitor selective for inhibition of Cox-2 and naproxen, an inhibitor of both Cox-1 and Cox-2, which differ in their propensity for GI adverse effects^8^. We analyzed the metabolome to seek discriminant biomarkers that might reflect and predict NSAID induced toxicity. Using both studies in humans and mice, we found that naproxen, but not celecoxib, depressed levels of tryptophan and its metabolites and that naproxen evoked fecal blood loss and cardiac inflammation in the mouse is rescued by tryptophan supplementation.

## Results

### Naproxen reduces tryptophan and kynurenine in humans

A double-blind three-way crossover study compared placebo, celecoxib and naproxen, each taken for 7 days. As expected, serum TxB_2_ and plasma PGE_2_, reflective of inhibition of Cox-1 and Cox-2 respectively *ex-vivo*, were both decreased by naproxen, while only PGE_2_ was reduced by celecoxib compared to placebo (Fig.1, A and B)^9,10^. The LC/MS/MS metabolomics data, obtained at T0 and T4, (i.e., 0hr and 4hr after final drug administration), were analyzed using Orthogonal Partial Least Squares-Discriminant analysis (OPLS-DA). The OPLS-DA scores plot (CV ANOVA of OPLS-DA plots for the T0 and the T4 time points are 0.0002, 0.0002 respectively) segregates the naproxen group from the celecoxib and placebo-treated groups (Fig. 1, D and E). We used the loading data from OPLS-DA analysis to identify metabolic pathways altered by naproxen treatment. The top metabolites corresponding to OPLS-DA between placebo and naproxen treatment were selected based on a Variable Importance in Projection (VIP) > 1.5 and were subjected to pathway analysis in Metaboanalyst. The pathway analysis shows that the false discovery rate (FDR) for tryptophan metabolism was 0.04, and the pathway impact was 0.2377 (Fig. 1F), consistent with this pathway being significantly altered by naproxen. Next, a paired 2-way ANOVA was used to perform univariate analyses on the metabolites obtained in the global metabolomics platform. This revealed significant changes in the levels of tryptophan and kynurenine in the plasma of naproxen-treated volunteers compared to those in the placebo and celecoxib groups (P < 0.01; Fig. 1, G and H). A targeted metabolomics experiment confirmed these findings and allowed us to obtain quantitative measures of the metabolites in the tryptophan pathway (Fig. 1I). The quantitative measurements of the metabolites demonstrate a decrease in the tryptophan levels at T0 and T4 in the naproxen group (P <0.0001, P<0.0001; fig S1A). The median (interquartile range) tryptophan level in plasma on naproxen was 68.0µmol/l (54.0 µmol/l, 82.7 µmol/l) while that on celecoxib it is 85.0µmol/l (70.5 µmol/l, 93.0 µmol/l) and on placebo is 91.1 µmol/l (88.6 µmol/l, 102.9 µmol/l) at T0. The median (interquartile range) of tryptophan on naproxen at T4 was 57.9 µmol/l (51.6 µmol/l, 63.8 µmol/l), and that for celecoxib, 83.4 µmol/l (75.6 µmol/l, 108.9 µmol/l) and for placebo, 86.0 µmol/l (79.3 µmol/l, 92.32 µmol/l) as shown in fig. S1A. The major tryptophan metabolite, kynurenine, was also depressed by naproxen at T0 and T4 (P <0.0001, P<0.0001; fig. S1A). The median level (interquartile range) of kynurenine on naproxen was 1.9 µmol/l (1.3 µmol/l, 2.9 µmol/l), on celecoxib, 3.1 µmol/l (2.0 µmol/l, 4.2 µmol/l) and on placebo, 3.0 µmol/l (2.3 µmol/l, 3.8 µmol/l) at T0. The concentrations of kynurenine on naproxen were 2.1 µmol/l (1.5 µmol/l, 2.4 µmol/l), for celecoxib 2.9 µmol/l (2.2 µmol/l, 3.4 µmol/l), and for placebo 3.2 µmol/l (2.2 µmol/l, 3.4 µmol/l) at T4 (fig. S1B). Other tryptophan metabolites, like kynurenic acid (KA), xanthurenic acid (XA), and anthranilic acid (AA), did not exhibit significant differences between the groups (fig. S1C-S1E). The Receiver Operating Characteristics curve (ROC) was used to evaluate the specificity and sensitivity of the marker molecules. The area under the curve (AUC) for tryptophan in placebo vs. naproxen was 0.84 (P<0.001; Fig.1J), celecoxib vs naproxen was 0.8 (P<0.001; Fig.1K), while placebo vs. celecoxib was 0.5 (P=0.23; Fig.1L). The AUC for kynurenine was 0.79 for placebo vs naproxen (P<0.001; Fig.1M); 0.78 for celecoxib vs naproxen (P<0.002; Fig.1N) and 0.5 for placebo vs. celecoxib (p=0.83; Fig.1O). Some of the other features discriminated at T0 vs T4 in naproxen treatment are nicotinamide and N-methyl-Nicotinamide along with sarcosine, arginine, proline, alanine, xanthine, and methyladenosine. Nicotinamide and N methyl nicotinamide also reflected alterations in tryptophan metabolism (fig. S1F, S1G and S1H). Isobutyrate, 3-hydroxybutyrate, 2-hydroxybutyrate, 2-oxoisocaproate, and succinate differed between T0 and T4 in the celecoxib group (FDR≤ 0.2; fig. S2). Apart from this, we also found alterations in the intermediates of the TCA cycle in urinary metabolic profiles from volunteers receiving both celecoxib and naproxen (fig. S3A). Consistent with these data, plasma alanine and urinary malonate are also altered by naproxen, reflecting an alteration in TCA cycle (fig. S3B-S3C). Additionally, pathway analysis obtained from the correlated urinary metabolites in celecoxib and naproxen-treated volunteers shows an alteration in the glycine serine and threonine metabolic pathway (fig. S3D).

**Fig. 1:**
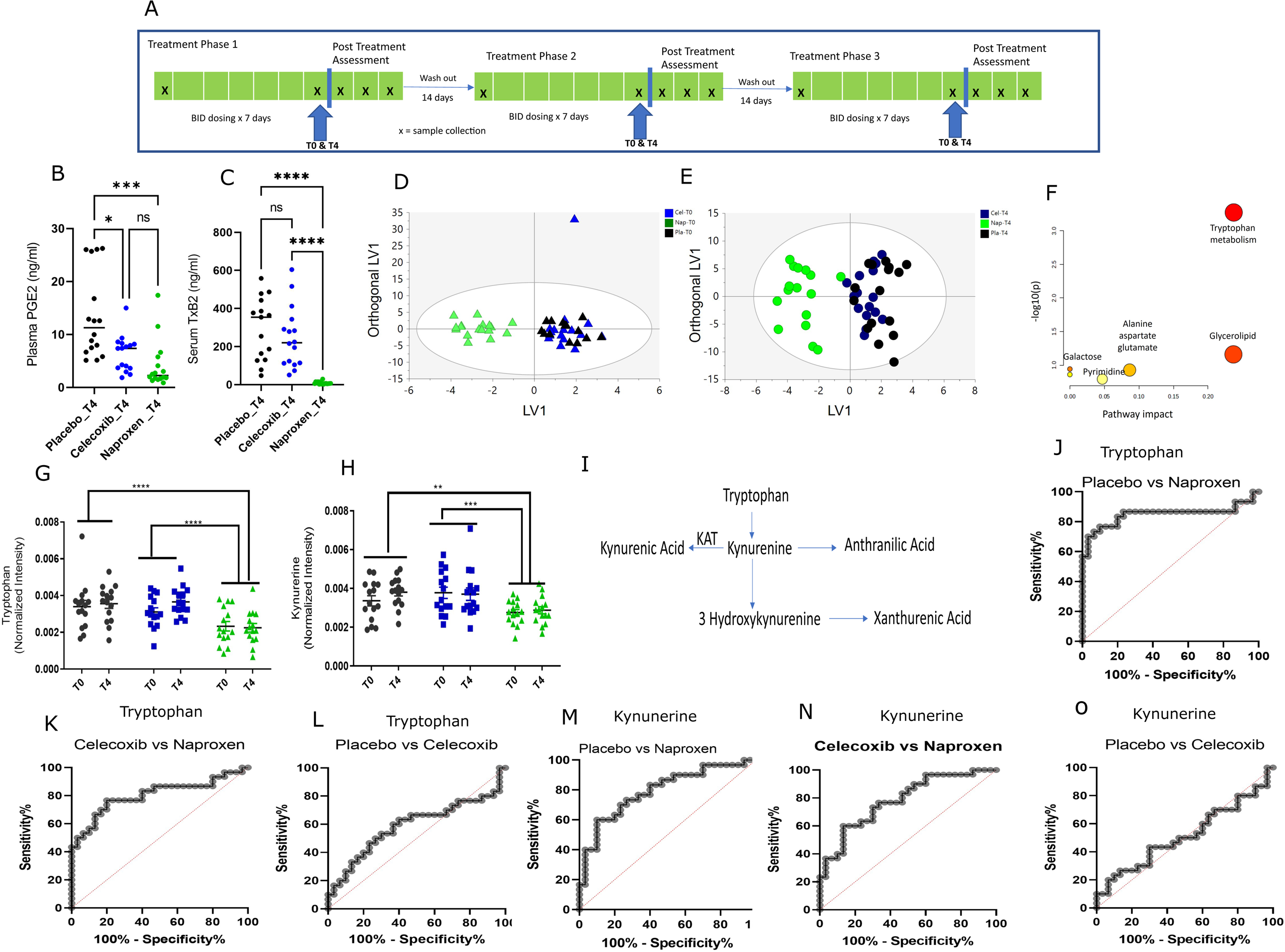
Metabolic profiles in volunteers in Naproxen alters wrt to Celecoxib and Placebo. A) The scheme for the clinical trial experiment for the human volunteers. B) Cox-1 inhibition *ex vivo* (Adapted from ref 8) and C) Cox-2 inhibition *ex vivo* by treatments (Adapted from ref 8) in a mixed effect analysis. D) OPLS-DA scores plot of plasma metabolites from Placebo, Celecoxib and Naproxen at T0. E) OPLS-DA scores plot of plasma metabolites from Placebo, Celecoxib and Naproxen at T4. F) Pathway analysis of the metabolites in global metabolomics platform. G) Univariate analysis of Tryptophan in Placebo, Celecoxib and Naproxen using two-way ANOVA. H) Univariate analysis of Kynurenine in Placebo, Celecoxib, and Naproxen using two-way ANOVA. I) The metabolic pathway of tryptophan metabolism. J-L) ROC plot of tryptophan in Placebo vs. Naproxen, Celecoxib vs. Naproxen, and Placebo vs. Celecoxib. M-O) ROC plot of Kynurenine in Placebo vs. Naproxen, Celecoxib vs. Naproxen and Placebo vs. Celecoxib. O). Black, blue, and green dots refer to control celecoxib and naproxen treatments. * designates statistical significance, *p ≤ 0.05, *** p≤ 0.001, **** p≤ 0.001.

### Naproxen reduced tryptophan and kynurenine in mice

Eight-week-old mice were randomized into naproxen and control groups (n=8/sex/diet), as shown in Fig.2A. After 3 weeks of dosing, naproxen had lowered tryptophan in the plasma compared to the controls (121.5 µmol/l; IQR: 106.8-133 µmol/l vs 147 µmol/l; IQR: 140.2-163.1 µmol/l respectively), (P<0.001; Fig.2B) in both sexes. Kynurenine was also reduced by ∼40% by naproxen (0.56 µmol/l; IQR: 0.63 µmol/l-0.42 µmol/l) compared to controls (0.99 µmol/l; IQR: 1.16 µmol/l-0.68 µmol/l), (P< 0.003; Fig.2C). The other metabolites in the tryptophan pathway, KA, XA, and AA, were not depressed by naproxen (Fig. 2, D, E, and F) consistent with our observations in humans. The AUC of the ROC was 0.91 for tryptophan, with a 95% confidence interval (CI) of (0.80 - 1.00), (P<0.001; Fig.2, G). The AUC for kynurenine was 0.85 with a 95% CI of (0.73-0.98), (P<0.0005; Fig.2H). Both tryptophan and kynurenine were also depressed by naproxen after only 5 days of treatment (fig. S4A-S4D). Weight and food intake were measured for three weeks to assess the potential effect of feeding on tryptophan levels. However, no significant differences were found between the groups (fig. S5A-S5E).

**Fig. 2:**
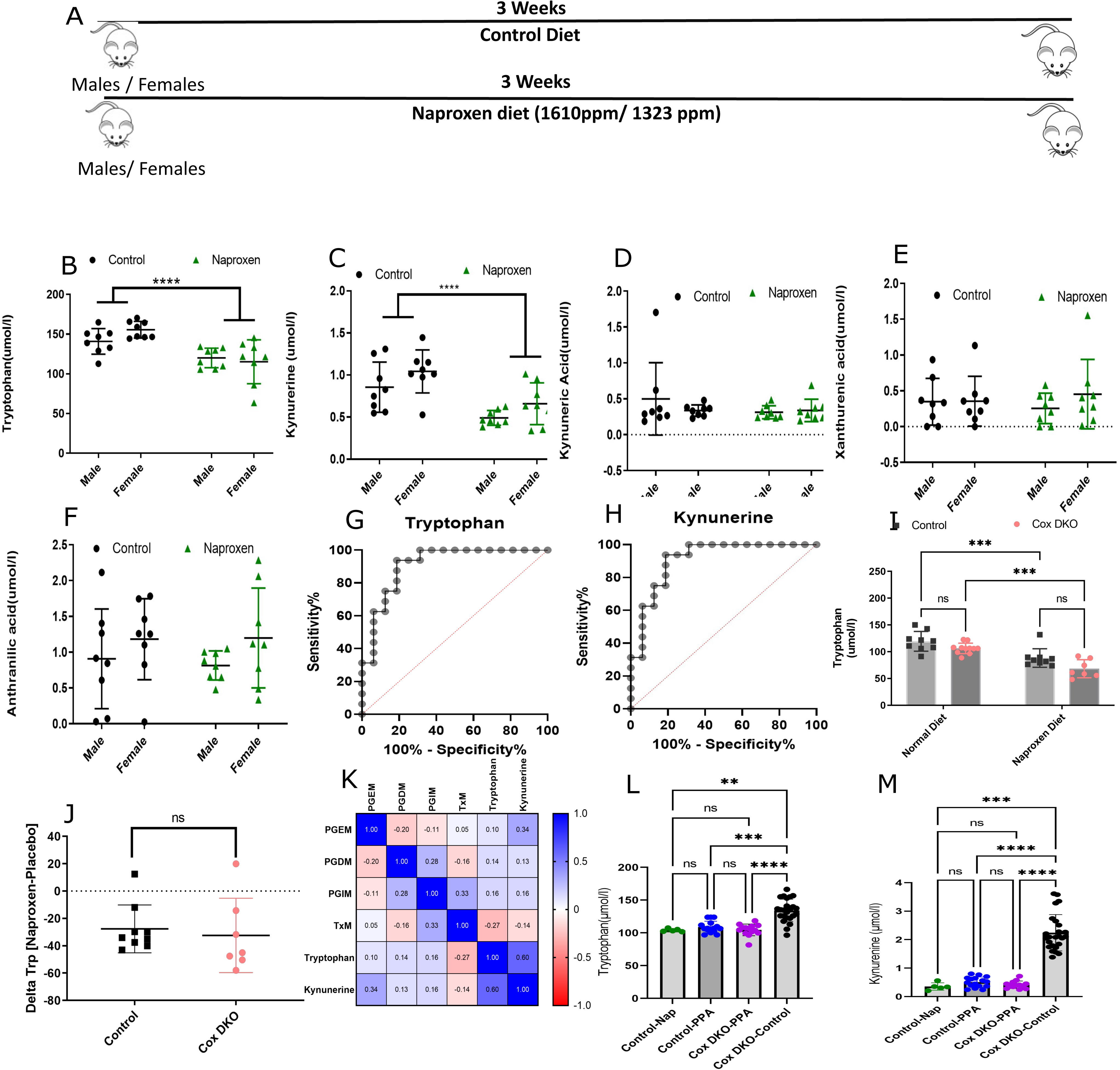
Plasma metabolic profiles of mice are altered by Naproxen diet. A) The scheme of the mouse experiment is to investigate the metabolic effects of Naproxen vs. the control diet. B) Targeted analysis of plasma tryptophan levels C) Plasma kynurenine levels. D) Plasma kynurenic acid. E) Plasma xanthurenic acid. F) Plasma anthranilic acid in mice by LC/MS/MS in a two-way ANOVA. G) ROC plot of Tryptophan in Control vs. Naproxen. H) ROC plot of Kynurenine in Control vs. Naproxen. I) Plasma tryptophan levels in Control mice and in *Cox*-DKO in control chow and Naproxen diet. J) Comparison of change in the tryptophan levels in control mice and *Cox*-DKO mice in Naproxen diet with respect to Control chow in a two-way ANOVA. K) Spearman correlation plot of human urinary prostaglandin metabolites and the plasma metabolite levels. L) Plasma Tryptophan levels of Control mice in Naproxen and R-2-Phenylpropionic acid diet (PPA) and “*Cox*-DKO” mice in PPA and control chow diet using One-Way-ANOVA. M) Plasma Kynurenine levels in Control mice in Naproxen and PPA diet and “*Cox*-DKO” mice in PPA and control chow diet using One-Way-ANOVA. The green, blue, purple and black circles denote the control mice in the Naproxen diet, control mice in the PPA diet, “*Cox*-DKO” mice in the PPA diet, and “*Cox*-DKO” mice in the control chow diet, respectively. All the data are expressed in mean ± SD * Designates statistical significance, **** p≤ 0.001.

To address the possibility that depression of tryptophan reflected platelet Cox-1 inhibition by naproxen, we measured plasma tryptophan and kynurenine in platelet-specific Cox-1 knockout (KO) mice (Pf4-Δ Cre ^+/-^/Cox-1^fx/fx^)^11^. Despite the suppression of the major Cox-1 product, Tx, in the KOs, tryptophan was unaltered, (fig.S6A-C). Similarly, we assessed the impact of naproxen on tryptophan in mice lacking both Cox-1 and Cox-2 globally, following postnatal depletion of Cox-1 and Cox-2 globally (*Cox* DKO) and controls, and there was no impact of genotype on drug effect (Fig. 2, I and J). Also, there was no significant correlation between prostaglandins with either tryptophan or kynurenine in naproxen-treated humans (Fig. 2, K).

We sought to address the possibility that tryptophan was differentially metabolized *via* the serotonin or melatonin pathways during naproxen administration. While melatonin was not detected in mouse plasma, consistent with previous findings^12^, the serotonin level was unaltered by naproxen (fig. S6D). We also found no evidence that naproxen altered kynurenine aminotransferase (KAT) activity as a potential cause for the change in plasma tryptophan concentration in both mice and humans (fig. S7A-D). R-2-phenylpropionic acid (PPA), a structural analog of naproxen, devoid of Cox inhibitory activity^13^ had no impact on tryptophan (Fig. 2, L and M).

### Naproxen alters tryptophan binding

Tryptophan was 73.8% bound to albumin in the placebo group (IQR: 69.90-78.66), while in the celecoxib group, it was 70.9 % (IQR:67.64 - 75.20) and in the naproxen group57% bound (IQR: 48.47-63.15). The %bound tryptophan was significantly decreased in naproxen compared to placebo and celecoxib (P<0.01 and P<0.05, respectively; Fig. 3A). Free tryptophan was reduced by naproxen in the plasma of volunteers (P<0.05; Fig.3B). Further, we found a negative correlation between tryptophan and naproxen levels in plasma (P<0.05; Fig.3C). We also found that bound tryptophan was decreased dose dependently by naproxen in mouse plasma (Fig.3D).

**Fig. 3:**
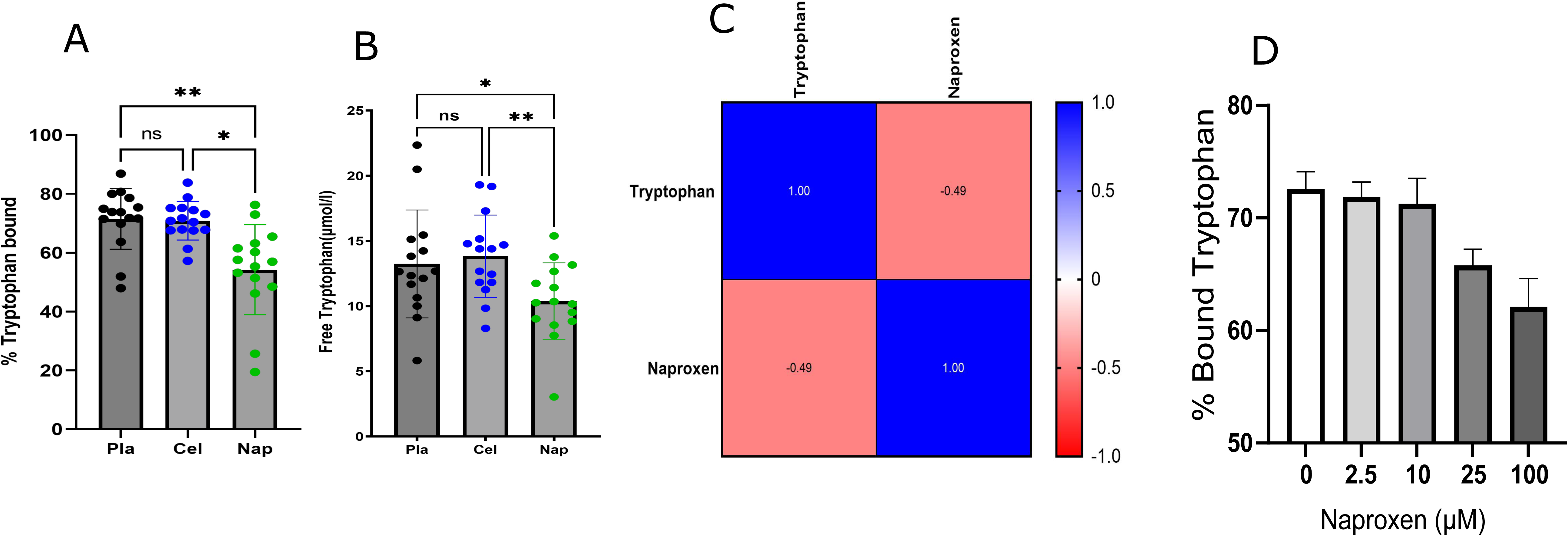
Naproxen displaces tryptophan from albumin binding. A) % bound tryptophan to albumin in plasma samples of volunteers in three different treatments: Placebo, Celecoxib and Naproxen analyzed in One-Way-ANOVA. B) Free tryptophan levels in Placebo, Celecoxib, and Naproxen in the buffer chamber of the RED plate analyzed by One-Way-ANOVA. C) Spearman correlation between plasma Naproxen and Tryptophan in the human volunteers. D) %Bound tryptophan concentration in mouse plasma treated with increasing doses of Naproxen. The data is expressed as mean ± sd. * Designates statistical significance, ** p≤ 0.01.

Importantly, both naproxen and tryptophan bind to the Sudlow 2 site of albumin with an association constant of 1.2–1.8 × 10^6^ M^−1^ and 4.88 × 10^4^ M^−1^, which explains the lower binding of tryptophan to albumin in the presence of naproxen. Our data are consistent with the displacement of tryptophan by naproxen.

### Potential role of naproxen-induced changes in the microbiome in tryptophan metabolism

Naproxen altered the microbiome, significantly increasing the *Coprococuccus* and *Ruminococcacae* taxa (FDR = 0.25, 0.25; Fig.4, A and B), both of which influence tryptophan metabolism^14,15^. This may explain the increase of urinary indole 3 acetic acid, and indole lactic acid in humans and mice (Fig. 4, E and F).

**Fig. 4:**
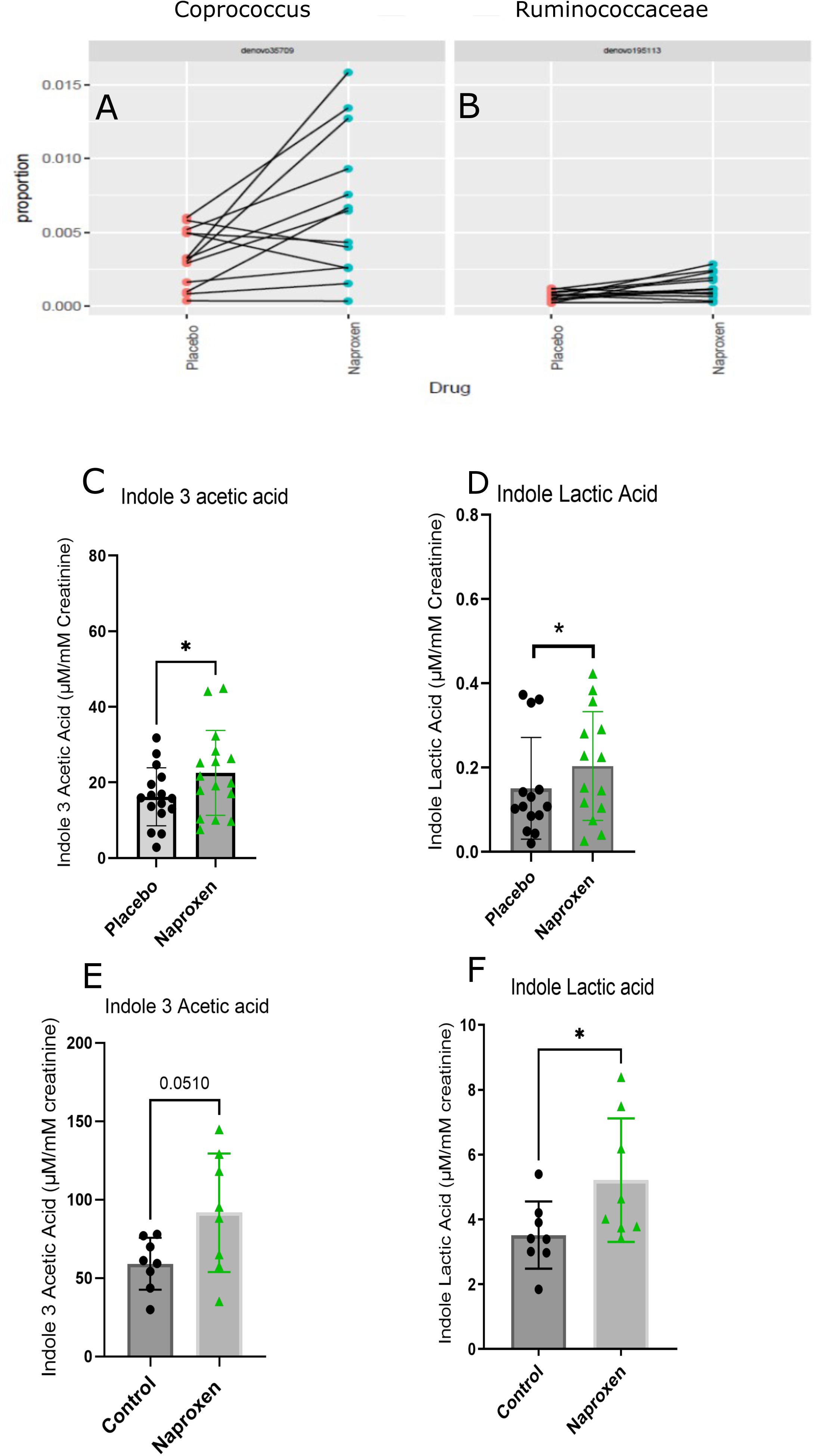
Naproxen alters microbiome and microbiome derived metabolites. A) Proportions of *Coprococcus* and *Ruminococcaceae* in human stool after Naproxen treatment. B) Urinary Indole 3 Acetic acid levels in Naproxen and Placebo treatment in human volunteers analyzed by paired Wilcox test. C) Urinary Indole 3 Acetic acid levels in Naproxen and Placebo treatment in mice by unpaired Welch test. The black and the green circles represent Placebo and Naproxen, respectively. The data is expressed as mean ± SD. * Designates statistical significance, * p≤ 0.05.

### Functional consequences of tryptophan depletion by naproxen

Depression of plasma tryptophan by NSAIDs was negatively correlated with a rise in Mean Arterial Pressure (MAP) in human subjects (https://clinicaltrials.gov/study/NCT02502006)^9^, suggesting a possible link between the change in tryptophan and cardiovascular function (fig. S8A). We supplemented mice on the naproxen diet with tryptophan (50mg/kg i.p. daily for 7 days) to address this possibility (Fig. 5A), as this was sufficient to restore plasma levels from their drug induced depression (Fig. 5B). While body weight was unaltered in these experiments, differentially expressed genes caused by naproxen were largely “rescued” by tryptophan supplementation (Fig. 5C). Of the 212 (out of 17093) genes that were rescued (Supplementary Table 1), 42% are involved in biological regulations, 45% in cellular processes, 24% response to stimulus and 18% in metabolic processes (PANTHER.db). Several inflammatory pathways are also restrained by tryptophan supplementation (Fig. 5D). An anti-inflammatory role of tryptophan in the heart is consistent with the negative z activation scores of “IL-13 activation pathway”, “cytokine storm signaling,” and “cardiac hypertrophy signaling,” (-0.447 -2.53 and -2.71, respectively), signifying inhibition of these pathways by tryptophan supplementation. Further, the causal network and upstream regulator analyses demonstrate that IL1β has a z activation score of -3.6 (BH corrected p-value = 0.000093), suggesting that IL1β might also be inhibited by tryptophan supplementation (Fig. 5E); this was confirmed by its measurement by RT PCR (Fig. 5F). “The nucleotide-binding domain, leucine-rich containing family, and pyrin domain containing 3” (NLRP3) was also decreased by tryptophan supplementation (Fig. 5G). Notably, NLRP3 activates Caspase-1, which in turn releases IL1β. Causal network analysis also revealed inflammasome inhibition by tryptophan supplementation (activation score of -4.12 (p=5.82e-8)). The genes rescued by tryptophan supplementation of naproxen treated mice which are also targets of IL1β are listed in Fig. 5H. These genes are involved in the “Inflammation-mediated chemokine and cytokine signaling pathway” and “Interleukin signaling pathway” (PANTHER.db, Fig. 5H). Additionally, a subset of IL1β mediated rescue genes is involved in various inflammatory pathways in the heart. These pathways are “cytokine storm signaling”, “IL-13 signaling”, “cardiac hypertrophy”, “cardiac dysfunction,” and “myocardial infarction,” demonstrated in Fig. 5I. Additionally, 1L-10 was rescued (fig.S9A), and IL6 receptor decreases following tryptophan supplementation compared to naproxen group (fig.S9B), suggesting a decrease in inflammation by the tryptophan supplementation. The elevation of IL17 receptor and IL4 receptor expression by naproxen was rescued by tryptophan supplementation (fig.S9C-D), both of which have been related to adverse cardiovascular health^16,17^, thereby consistent with a beneficial role of tryptophan in this context. Furthermore, tryptophan also plays a role in mitigating energy demand in heart by rescuing FNIP2, PGC1α, ITGB3 and TGM2. As shown by previous studies, these genes play a crucial role in energy metabolism and mitochondrial function^18–21^. In addition to this cardiovascular impact of tryptophan supplementation, it also prevented the increase in fecal hemoglobin caused by naproxen (Fig S10).

**Fig. 5:**
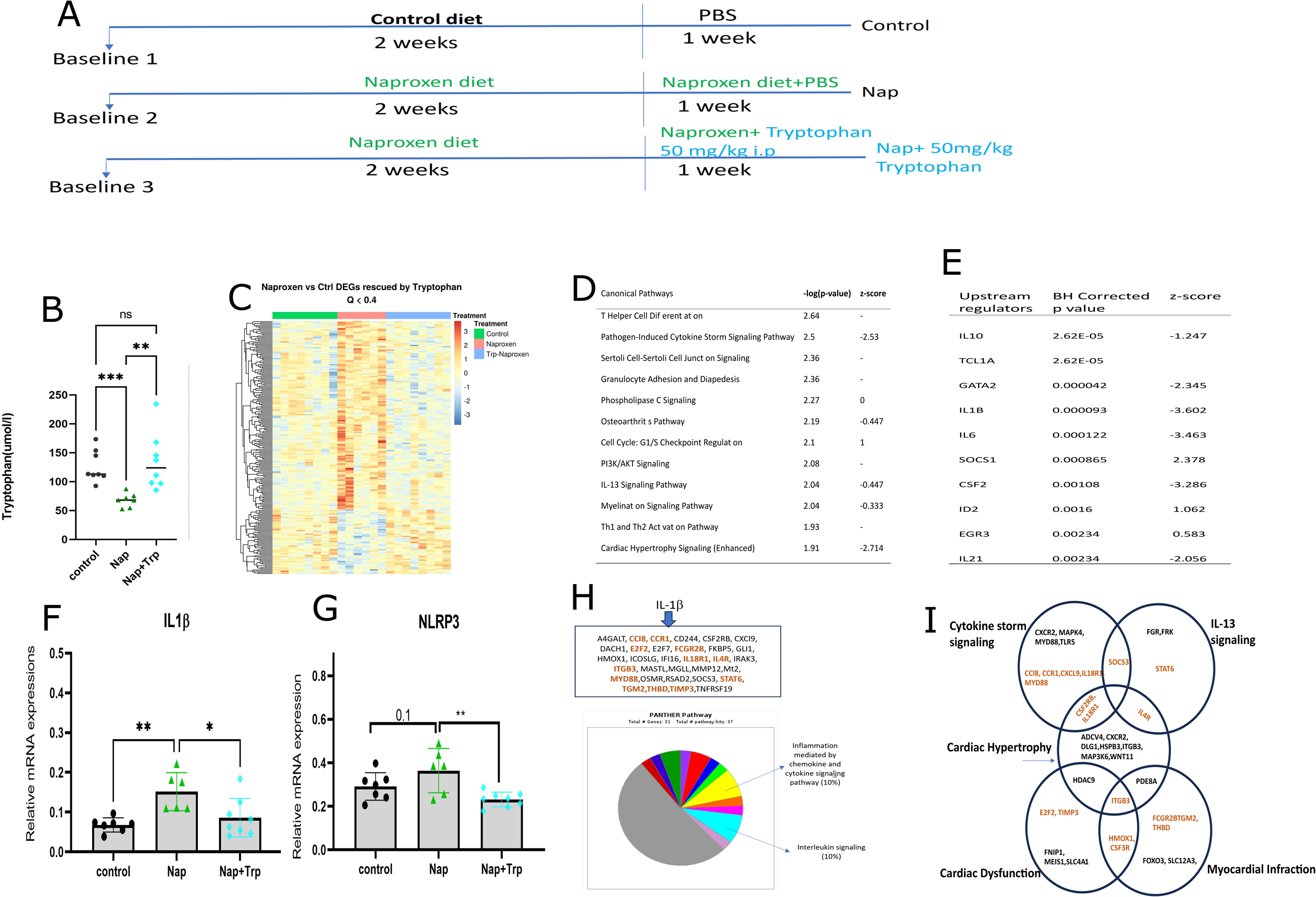
Tryptophan Supplementation experiment in mice during Naproxen treatment ameliorate inflammation. A) Scheme of Tryptophan supplementation experiment in mice. B) Plasma Tryptophan levels in three different groups (Control, Naproxen, Naproxen+ Tryptophan). C) Heatmap showing “rescue genes” in the heart after the supplementation of tryptophan with Naproxen. D) Canonical Pathway analysis for the rescue genes in Ingenuity Pathway Analysis (IPA). E) Upstream regulators of the rescue genes in IPA. F) IL-1β levels in the mouse heart after tryptophan supplementation in Naproxen treatment analyzed by One-Way-ANOVA. G) NLRP3 levels in mouse heart following tryptophan supplementation in Naproxen treatment One-Way-ANOVA. H) Target genes of IL-1β that are rescued by tryptophan in Naproxen treatment. Pathway analysis of the target genes. I)Venn diagram showing the target genes of IL-1β that are part of possible cardiac phenotypes in mice in Naproxen treatment in IPA analysis. Black circles, green triangles, and blue diamonds refer to control, naproxen, and naproxen+tryptophan groups. The data is expressed as mean ± sd. * designates statistical significance, * p≤ 0.05, ** p≤ 0.01 respectively.

## Discussion

NSAIDs are amongst the most consumed drugs because of their non-addictive efficacy in relieving pain and inflammation. Despite this, they can cause gastrointestinal and cardiovascular adverse effects which can, on occasion, be life threatening^22^. While metabolomic analyses of the response to NSAIDs have been reported^23^, here we study the comparative effects of a traditional NSAID, naproxen and one specifically designed to target Cox-2, celecoxib^24^ in humans. Gastroduodenal adverse effects of NSAIDs have been attributed largely to the inhibition of COX-1^25^, while their risk of heart attack, hypertension and stroke reflects particularly suppression of Cox-2^26^.

Our most striking finding was that naproxen, but not celecoxib, depressed tryptophan and its major metabolite, kynurenine. We replicated this drug effect in mice. A surprise was that this seems unrelated to suppression of PGs and Tx by naproxen. Using a variety of mutant mice^11,27,28^, we found that depletion of Cox-1 or Cox-2, alone or together had no impact on the depression of tryptophan by naproxen, uncoupling this effect from drug induced suppression of PGs and Tx.

The next question was how naproxen might depress tryptophan. Although some effects on other metabolites – elements of the TCA cycle, for example – were observed, the impact on the tryptophan pathway was the most striking. Measurement of free and bound tryptophan provided us with evidence consistent with naproxen, but not celecoxib, displacing tryptophan from its protein binding site, permitting its degradation and elimination under steady state dosing conditions. Indeed, this may interact with perturbation of the microbiome by naproxen to accelerate tryptophan disposition. Thus, we found that the drug increased the abundance two fecal strains, *Ruminococcaceae* and *Coprococcuses*, which have been shown to influence tryptophan metabolism^29–32^ by its conversion to indole derivatives. The next question was whether this depressive effect of naproxen on tryptophan might have contributed to its adverse effect profile. To address this possibility, we examined the impact of tryptophan on gene expression in mouse heart. Here, using a variety of approaches, we found that naproxen induced expression of inflammatory genes, including those driven by IL-1β, culminating in activation of the inflammasome. Supplementation of these naproxen treated mice with tryptophan, sufficient to restore its plasma levels rescued much of the inflammatory signature of drug induced gene expression.

Correspondingly, we performed a preliminary study to determine if this mitigating effect of tryptophan supplementation might influence other aspects of the adverse effect profile of naproxen. Naproxen increases fecal occult blood, reflective of the gastroenteropathy it causes in mice and humans. Here again, tryptophan supplementation appeared to attenuate the damage.

In summary, we report that naproxen, in humans and mice, depresses tryptophan and its major metabolite by a mechanism unrelated to COX inhibition and reflecting displacement from albumin and accelerated disposition by the gut microbiome. Naproxen induces an inflammatory profile in mouse heart that may contribute to the heart failure that complicates NSAID administration. This signature and naproxen induced fecal blood loss are both rescued by tryptophan supplementation. Thus, depletion of tryptophan may contribute to the cardiovascular and gastrointestinal adverse effects of this and perhaps other NSAIDs in humans. Our studies in mice raise the possibility that tryptophan supplementation my attenuate adverse effects and widen the therapeutic index of this commonly consumed NSAID.

## Methods

### Study Design of Clinical Trial

Healthy volunteers (N=15; 9 men, 6 women) were treated with naproxen (250 mg bid), celecoxib (100 mg bid), and placebo for seven days in a double-blind, randomized, three-way cross-over design (https://clinicaltrials.gov/study/NCT02502006)^9^. There was a washout period of at least 14 days in between the arms. The participants were asked to abstain from analgesics until study completion. All the experiments were performed in a blinded manner by the analyst. The trial design is shown in (Fig. 1A).

### Metabolomics analysis in plasma

Plasma samples taken at T0 (T=0h after 7 days of dosing) and T4 (T=4h after 7 days of dosing) were subjected to metabolomics analysis. The time points represent the crest and the trough of the pharmacological activity of the drugs. Semi-targeted mass spectroscopy was employed as an initial screen. Polar metabolites were extracted from 50µL of plasma using a modified Bligh Dyer protocol^33^. Quality control samples were prepared by pooling all the samples and were used for run order corrections. MS/MS analysis was performed using a Waters micro TQS (Waters Corp., Milford, MA.), with electrospray ionization. Data were integrated into Waters Targetlynx 4.1 software (Waters Corp., Milford, MA.) and further processed in R. Every metabolite feature was fitted using a locally weighted scatterplot smoothening function. Additionally, probabilistic quotient normalization was applied to the data. The multivariate analysis of the data was performed in Simca P+ (Umetrics Inc., Umea, Sweden).

### Measurements of prostanoids in the ex-vivo assay

The ex *vivo* measurements of Cox-1 and Cox-2 activity was performed as described in the literature previously^9^. LC/MS/MS was used to quantify prostanoids as described^34^. Briefly, 5ng each of d^4^ PGE_2_ and d^4^ TXB_2_ were added to 100 µl of plasma and 50µl of serum, respectively. This was subjected to solid-phase extraction (Strata-X, 33µm Polymeric Reversed Phase, Phenomenex) and measured by LC/MS/MS ^35,36^ using a Waters AQUITY UPLC system. Data were integrated into Targetlynx 4.1 software (Waters Corp., Milford, MA.).

### Targeted Metabolomics of Tryptophan Metabolism Pathway

For the targeted assay of the metabolites, 40µl of the plasma samples were processed as previously described^37^. 10µl of an internal standard, consisting of 260 ng of d^5^-Tryptophan and 0.1% formic acid, was added to the solution. This was followed by Solid Phase extraction in the reversed-phase cartridge (Strata-X, 33µm Polymeric Reversed Phase, Phenomenex, Torrance, CA). The samples are dried and reconstituted in 40µL acetonitrile and 360µL water. Separation of the metabolites was performed using an ultra-performance liquid chromatography (UPLC) column, 2.1 × 150 mm with 1.7 μm particles (Waters ACQUITY UPLC BEH C18) following this gradient: Mobile phase A consisted of 0.1% formic acid 95% water 5%B, and mobile phase B was acetonitrile: methanol in 95:5 containing 0.1% formic acid. Linear gradient was run as follows: 0 min 98% A; 5 min 95% A; 5.5 min 90% A; 20 min 40% B; 20.1 min 90% A; 23 min 98% A at a flow rate of 0.3 mL min^−1^ with a total run time of 25 min. Calibration curves were used for the quantitation of the metabolites. Data were integrated into Waters Targetlynx 4.1 software.

### Measurement of plasma drug concentrations in blood

For the analysis of drug concentration, 50µl of an internal standard containing 500 ng of d^3^ Naproxen was added to plasma samples before solid-phase extraction (Strata-X, 33µm Polymeric Reversed Phase, Phenomenex). A Waters Triple Quadrupole Mass Spectrometer equipped with a UPLC system was used to measure drug concentrations using a 2.1 × 150 mm column, with 1.7 μm particles (Waters ACQUITY UPLC BEH C18) following this gradient: the mobile phase consisted of 0.5% ammonium acetate at pH 5.7 (mobile phase A) and acetonitrile (mobile phase B). The following gradient was then used with a flow rate 350 µl/min; (0-20 min 20% B-90% B; 20-22 mins 90-100% B, 22-30 mins 100-20%B). Data were again integrated into Targetlynx 4.1 software.

### Measurement of bound and free tryptophan concentrations in blood

Free and bound tryptophan were separated by equilibrium dialysis (RED plate, Thermo Fisher). Briefly, 100µl of plasma was inserted into the inner vial and 300µl of the buffer (provided in the kit) was added in the outer vial. The plate was shaken for 4 hrs. The samples were removed from the chambers and stored in separate vials at -80^0^C. Before the LC/MS/MS analysis, the samples were thawed in ice and d^5^-tryptophan internal standard was added. To understand the direct role of naproxen on tryptophan displacement, pooled mouse plasma was aliquoted to 10 equal parts of 100µL. Duplicate samples were treated with (0µM, 2.5 µM,10 µM, 25 µM, 100 µM) of Naproxen *in vitro* and were subjected to Equilibrium Dialysis in RED plate as mentioned before. The samples were further processed for the tryptophan measurement.

### Analysis of Indole Metabolites in urine

Urine was collected from mice and humans for analysis of Indole 3 Acetic acid, Indole lactic acid, and Indole sulfate by LC/MS/MS. 10µl of urine sample was added to 100µl of milli-Q water containing d^5^-Tryptophan as internal standard. The mixture was centrifuged at 10,000g at 4^0^C for 10 mins. The supernatant was loaded to an autosampler, and separation of the metabolites was achieved by using a Waters ACQUITY UPLC system as previously described using a mobile phase A of 95:5 water and B, with 0.1% formic acid, and mobile phase B consisting of 95:5 acetonitrile: water with 0.1% formic acid. The metabolites were normalized with respect to urinary creatinine, also quantified by LC/MS/MS as previously described.

### Global Metabolomics profile of urine by Nuclear Magnetic Resonance

Urine samples were centrifuged at 13300 rpm at 4^0^C. The samples were aliquoted, and 180 µL of the aliquot was added to 20 µL 1M phosphate buffer containing 2.5 M DSS and 0.03%(v/v) sodium azide. The samples were transferred to 3 mm NMR tubes (Bruker Biospin, Billerica, MA). ^1^H NMR spectra were acquired in a 700 MHz Bruker Avance III HD NMR spectrometer (Bruker Biospin, Billerica, MA) fitted with a 3mm triple resonance inverse (TXI) probe. All spectra were acquired using a NOESYPR1d pulse program with relaxation delay of 1s, 0.1s mixing time, 76k data points and 14ppm spectral width. A total of 256 scans were acquired per sample. Water was suppressed using the presaturation technique during relaxation delay and mixing time. Raw spectral data were imported into Chenomx v8.0 (Chenomx Inc. Edmonton, Alberta, Canada) for further processing. The spectra were Fourier transformed after zero filling to 128k, and Raw spectral data were imported into Chenomx v8.0 (Chenomx Inc. Edmonton, Alberta, Canada) for further processing. Fourier transformed the spectra after zero filling to 128k, and linear broadening of 0.1 Hz was applied. All spectra were referenced to internal standard followed by targeted profiling of metabolites of interest^38^. The peaks of the spectra are profiled using Chenomx software.

### Calculation of sample size for mice experiment

We determined the sample size for the animal experiment using online software (clincalc.com/stats/samplesize.aspx). Considering the human trial outcome of tryptophan concentrations reflecting effect size (Cohen’s d) to be 0.94 between placebo and naproxen treatment, we aimed to minimize type II error and type I error set at α=0.05 and sampling power at 80%, a total of 12 samples were needed in each group. However, considering a dropout of mice due to chronic naproxen dosing during experiment, we decided to start with N=16 in each group.

### Animal experiments to study tryptophan metabolism on Naproxen

Male and Female C57BL/6 mice were obtained from the Jackson Laboratory (Bar Harbor, ME) at six weeks of age. After two weeks of acclimatization, they were randomly assigned to two groups, n=16 per group: Control Chow (Male=8, Female=8) and Naproxen diet (Male=8, Female=8). Animals were kept in a 12-hour day/night cycle and had ad libitum access to food and water. The food was provided as pellets (Naproxen diet, 5001; 1323 ppm for females and 1610 ppm for males). Weights were recorded weekly for all the mice for three weeks. Animals were sacrificed at the end of three weeks. Blood was collected, and plasma was immediately separated after centrifugation at 3000g for 5 minutes (Eppendorf Centrifuge 5424). The plasma obtained was frozen at -80^0^C for analyses. Platelet specific Cox-1 KO mice (Pf4-Δ Cre ^+/-^/Cox-1^fx^/^fx^) were generated as mentioned before^11^. The Cox-1^fx/fx^ mice were kindly provided by Harvey Herschman at UCLA^39^. For global Cox-1 deficient mice, Ind-Cre**^+/-^** mice^40^ were mated with Cox-1^fx/fx^ mice to generate the Ind-Cre**^+/-^** Cox-1^fx/fx^ mice. Ind-Cre**^+/-^**Cox-2^fx/fx^ were generated as previously described ^27^. To avoid the roles of Coxs during development we generated mice in which both Cox-1 and Cox-2 were depleted postnatally in a tamoxifen inducible manner ^28^. These Cox-1^fx/fx^ Cox-2^fx/fx^ CMV-Cre^+/-^ mice are abbreviated as “*Cox*-DKO” mice in this paper.

### COX enzymes and tryptophan metabolism in mice on Naproxen

We utilized mice in which Cox enzymes were depleted to address the role of their inhibition in mediating the effects of Naproxen on tryptophan. “*Cox*-DKO” and their controls were profiled for their plasma tryptophan response to naproxen administration. In a different experiment, control mice were treated with Phenyl propionic acid (PPA) in the diet (1323 ppm R2 Phenyl propionic acid) and were compared to control mice on naproxen (1323 ppm). The mice were allowed to feed *ad libitum* for 3 weeks prior to plasma collection.

### Hemoglobin and Calprotectin measurements

Human and mouse fecal samples were assayed for hemoglobin and calprotectin using ELISA kits from MyBioSource (San Diego, CA) and Alpco (Macedon, NY) respectively. The samples were processed according to the manufacturers’ instructions.

### Analysis of the human fecal microbiome

Analysis using 16S amplicon sequencing was carried out as described^41–43^. Briefly, DNA was isolated from ∼200 mg of stool using the Qiagen PowerSoil Kit (Lot #154048139) following the manufacturer’s protocol. Isolated DNA was quantified using the Picogreen method. Primers to amplify the bacterial 16S rRNA gene region were barcoded to label each sample and PCR reactions were carried out in triplicate using Accuprime (Invitrogen, Carlsbad, CA, USA). Each reaction contained 5ul of extracted DNA and 0.4uM of each primer 50 nanograms of DNA and 10 pM of each primer. Primers annealing to the V1V2 region of the 16S bacterial gene were used for amplification as described ^44^. Amplified 16S rDNA was purified using a 1:1 volume of Agencourt AmPure XP beads (Beckman-Colter, Brea, CA, USA). The purified products were pooled in equal amounts and analyzed using Illumina MiSeq sequencing. DNA free water and blank extraction columns were subjected to the same purification and amplification procedure to allow empirical assessment of environmental and reagent contamination. Positive controls were also included, consisting of synthetic DNA plasmids mimicking sequences from well-studied organisms. Quality control and sample analysis was essentially as described in ^41^.

### Tryptophan rescue experiment in mice

Female C57BL/6 mice (N=24; N=8 per group) from Jackson Laboratory (Bar Harbor, ME) were grouped and treated as “control”, “naproxen,” and “naproxen + tryptophan” for three weeks of the study. The mice in control group were on the normal chow while that in “naproxen” and “naproxen + tryptophan” group received the Naproxen diet (Naproxen diet, 5001; 1323 ppm). The mice in the rescue group received intraperitoneal (i.p.) injection of 50mg/kg Tryptophan in Phosphate buffer Saline (PBS) for 1 week as shown in Fig 5A. All mice on the “naproxen” and “control” group received i.p. injection of PBS for 1 week.

#### RNA-Seq Quantification

The RNA was extracted from heart using a Promega kit (AS1340). RNA samples were quantified with NanoDrop 8000 Spectrophotometer and Agilent 21200 Bioanalyzer systems. The samples were run utilizing an Illumina Truseq mRNA RNAseq library prep kit. The samples were run utilizing with 400 ng of the input. After library prep and QC using Tapestation they were normalized to 1.55nM and run on the Illumina NovaSeq 6000 sequencer using an v1.5 S2 200 cycle kit aiming for 40 million reads per sample. Further, we used Salmon (v1.9.0)^45^ to quantify transcript read counts from the raw FASTQ files against transcripts from v102 of the Ensembl annotation^45^. We conducted these analyses across two RNA-Seq experiments (Experiment 1: mix of male and female mice treated with vehicle or naproxen. Experiment 2: female mice treated with vehicle, naproxen, or tryptophan + naproxen). The Salmon index we used for these data includes transcript sequences from the ‘Mus_musculus.GRCm38.cdna.all.fa’ (all Ensembl transcripts, excluding non-coding RNAs) and ‘Mus musculus.GRCm38.ncrna.fa’ (non-coding RNAs) FASTA files, both of which we downloaded from the Ensembl ftp site. We also included the full DNA sequence from the primary assembly (GRCm38; also downloaded from Ensembl) in the Salmon index as a decoy. When running Salmon, we used the following command line parameters to account for sequence, GC, and positional biases: “-- seqBias --gcBias –posBias”. Next, we imported the Salmon output into R (v4.2.2) and summarized these data to gene level quantifications with the tximeta package (v1.16.0)^46^. Unless otherwise indicated, all analyses and visualizations in this manuscript were prepared from the unnormalized, gene-level read counts estimated by Salmon and tximeta.

### Differential Gene Expression

We used DESeq2 (v1.38.1)^47^ to conduct differential expression (DE) analyses between our three groups of interest (control, naproxen-treated, naproxen + tryptophan-treated). For the ‘control vs naproxen’ DE comparison, we used data from both RNA-Seq experiments. When fitting the model for the ‘control vs naproxen’ experiment, we included a term to account for any batch effects between the two experiments. We only used data from the second experiment for the ‘control vs. naproxen + tryptophan’ and ‘naproxen vs. naproxen + tryptophan’ DE analyses because the first experiment did not include any mice treated with tryptophan. For all comparisons, we allowed DESeq2 to apply its default minimum expression filter, outlier detection, and Benjamini-Hochberg multiple testing adjustment to p-values (also known as q-values). Next, we merged the results of these DE analyses to identify genes affected by naproxen treatment, where those effects were reversed (or ‘rescued’) by tryptophan treatment. Briefly, we identified genes with evidence of DE (q-value < 0.4) in both the ‘control vs naproxen’ and ‘naproxen’ vs ‘naproxen + tryptophan’ comparisons. We further reduced this list to those with fold-change values showing opposite signs in these two comparisons. In other words, we identified DE genes that showed a simultaneous increase in response to naproxen treatment and a decrease in response to tryptophan supplementation (or *vice versa*).

### Pathway Analyses

We conducted all pathway analyses with the QIAGEN Ingenuity Pathway Analysis tool (IPA). Unless otherwise stated, we used the list of all provided genes as the background gene set for enrichment analysis instead of the list of genes contained in the Ingenuity Pathway Analysis knowledge base.

### Quantitative Real-time PCR of genes

The RNA was extracted using a Promega kit (AS1340). RNA samples were quantified with NanoDrop 8000 Spectrophotometer and Agilent 21200 Bioanalyzer systems. Quantitative Real-time PCR was performed using Taqman Gene Expression Assays for IL-1β (MM00434228_m1), IL-6, IL-10, IL-10RA.

### Statistical analysis

The metabolomics data set from plasma and urine from human volunteers was analyzed in R 4.1.2. Statistical testing was reported after correction for multiple testing using the Benjamini Hochberg (BH) method. Two-way analysis of variance (ANOVA) was applied to the targeted analysis of metabolites in Prism 9, followed by a pairwise test as appropriate. Mann Whitney test, Wilcoxon test analysis was conducted as indicated in figure legend. *designates statistical significance, *p ≤ 0.05, **p≤ 0.01, *** p≤ 0.001, **** p≤ 0.001.

## Acknowledgments

Dr. FitzGerald is the McNeil Professor of Translational Medicine and Therapeutics.

## Disclosures

Dr. FitzGerald is Senior Advisor to Calico Laboratories.

We acknowledge the technical support of Wang, Fengxiang. 16Ssequencing was provided by the CHOP Microbiome Center, Children’s Hospital of Philadelphia.

S.G. has nothing to disclose.

## Authors Contributions

SG, KB, SYT, KT, ER, and RJ conducted the experiment. SG generated data and figures interpreted the results and wrote the manuscript. AS and NL analyzed data and wrote manuscript. FDG interpreted the results. TG and GAF planned the project, supervised the study, interpreted the results, edited the manuscript and acquired funding.

## Supplementary Figure legends

**fig. S1:**
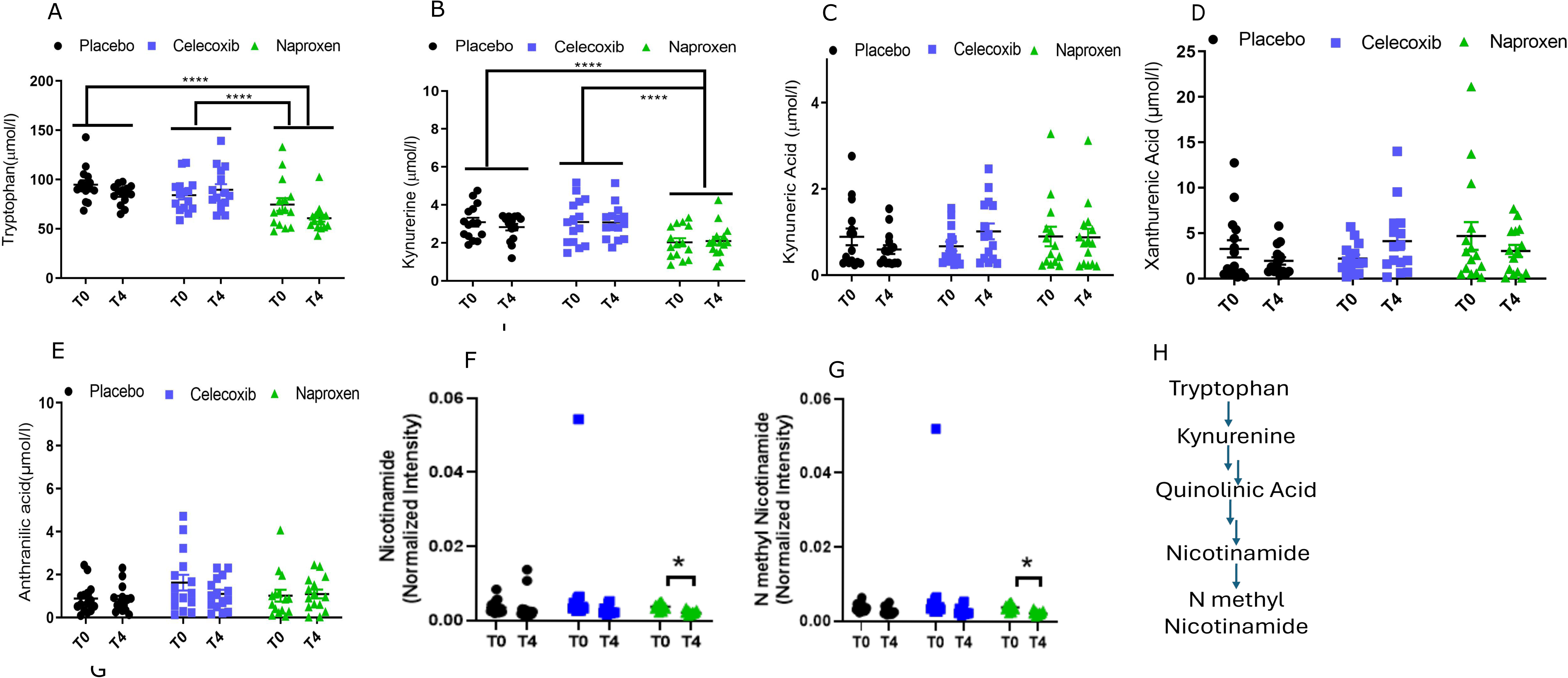
Targeted metabolite profiling of plasma tryptophan metabolites in human volunteers in a clinical trial validates global metabolic profiling. A) Tryptophan B) Kynurenine C) Kynurenic Acid D) Xanthurenic acid and E) Anthranilic acid by LC/MS/MS using two-way ANOVA. F) Nicotinamide by paired Wilcox test. G) N Methyl Nicotinamide by paired Wilcox test. The black, blue, and green circles indicate placebo, celecoxib, and naproxen. * designates statistical significance, * p≤ 0.05, **** p≤ 0.0001 respectively.

**fig. S2:**
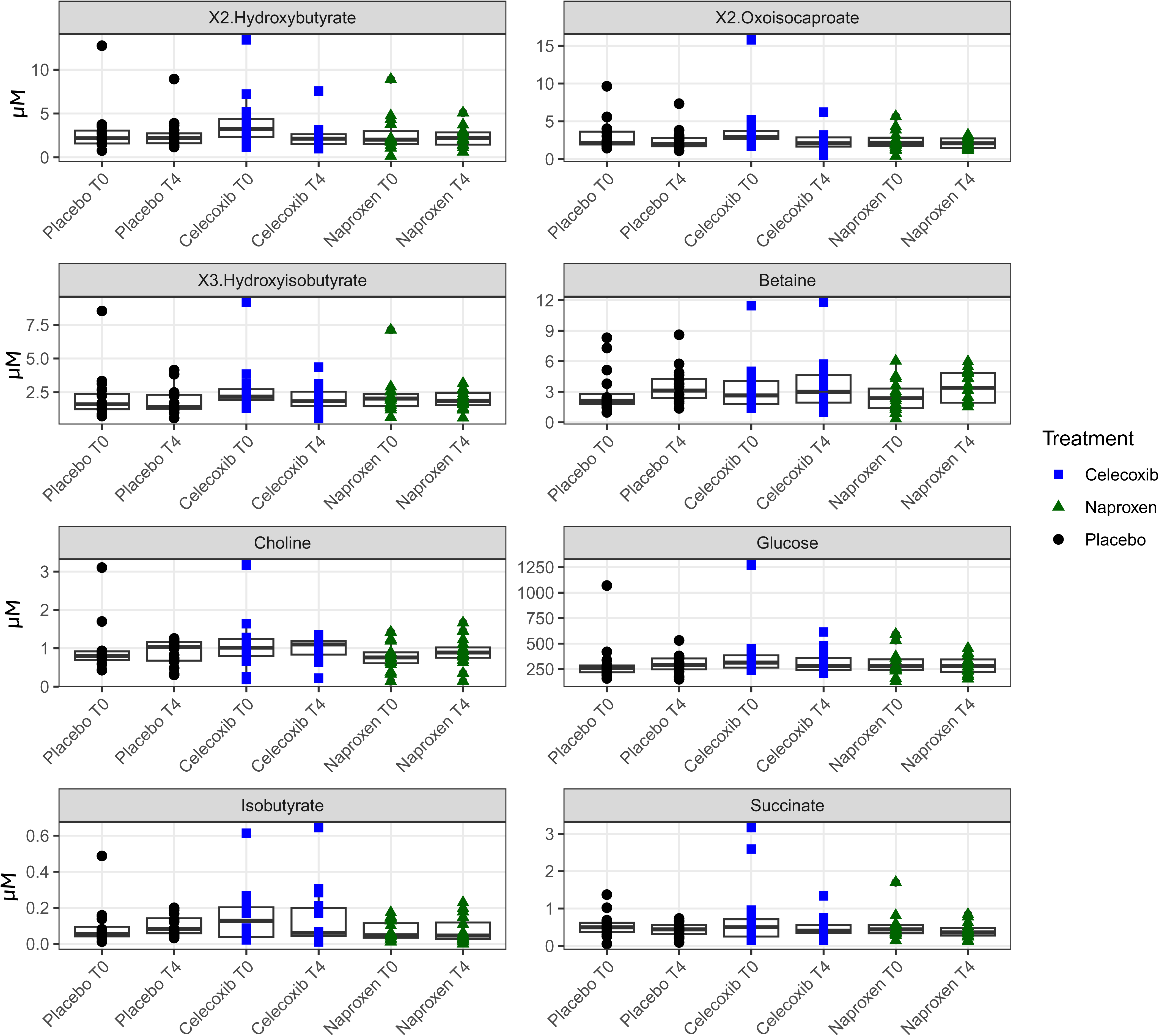
Plasma metabolites altered in Celecoxib treatment with respect to Placebo and Naproxen (FDR ≤ 0.02) suggest changes in amino acid metabolism. **Metabolic profiling of** A)2-Hydroxybutyrate B)2 Oxoisocaproate C)3 Hydroxyisobutyrate D) Isobutyrate E) Succinate F) Betaine G) Choline H) Glucose profiled by ^1^H NMR spectroscopy in plasma of human volunteers in placebo, celecoxib and tryptophan treatment. The data is expressed as mean ± SD.

**fig. S3:**
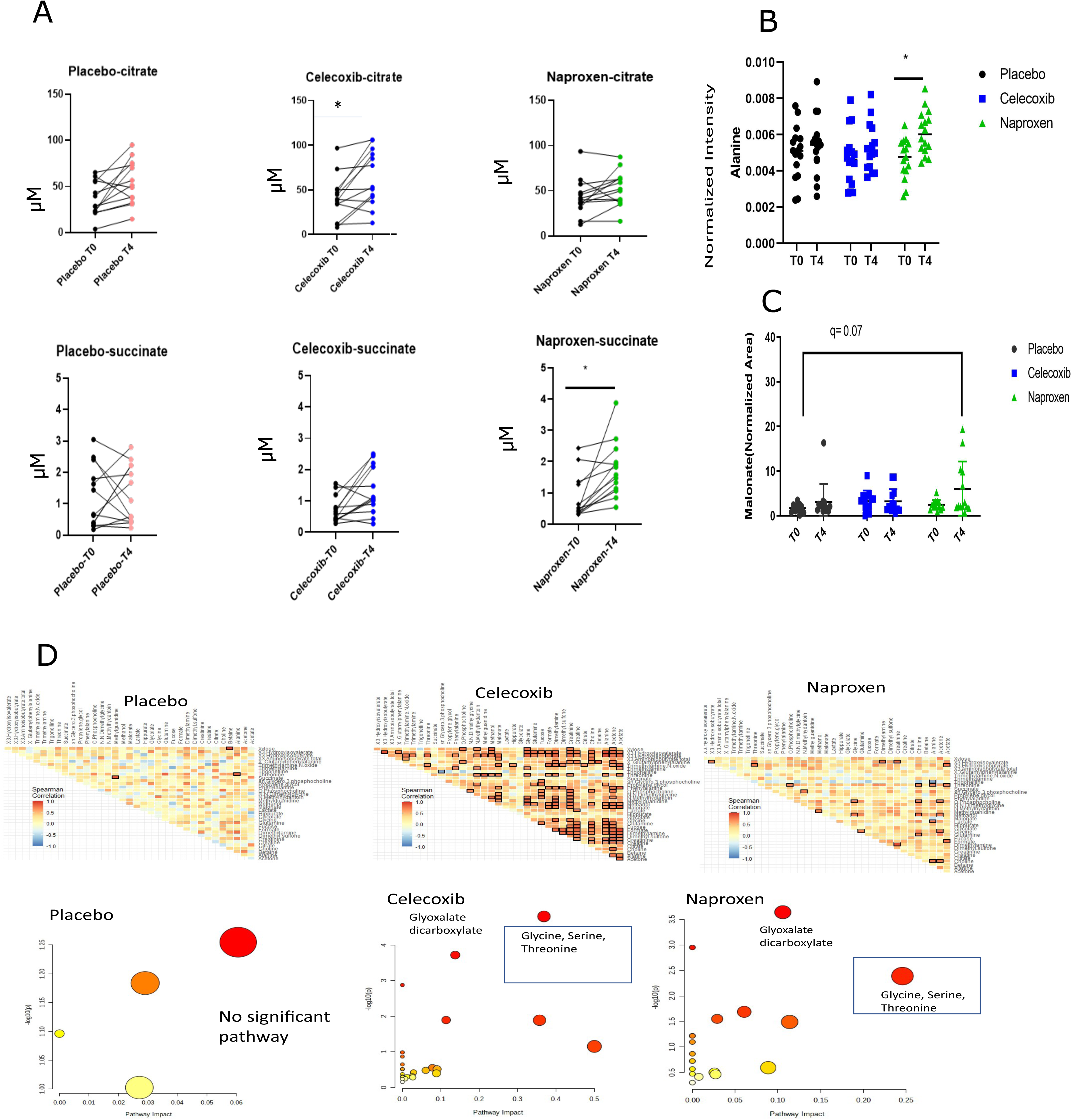
NSAID administration alters the TCA cycle and amino acid metabolism. A) Urinary levels of citrate and succinate in paired groups of Placebo-T0 vs. Placebo T4, Celecoxib-T0 vs. Celecoxib-T4, Naproxen-T0 vs. Naproxen-T4. B) Plasma alanine in human volunteers in Placebo, celecoxib and naproxen treatment. C) Urinary malonate human volunteers in Placebo, celecoxib and naproxen treatment. Black circles, blue squares and green triangles refer to control, celecoxib, and naproxen treatment. D) Spearman correlation analysis of urinary metabolites in human volunteers and Pathway analysis of the correlated metabolites. * designates statistical significance, * p≤ 0.05.

**fig. S4:**
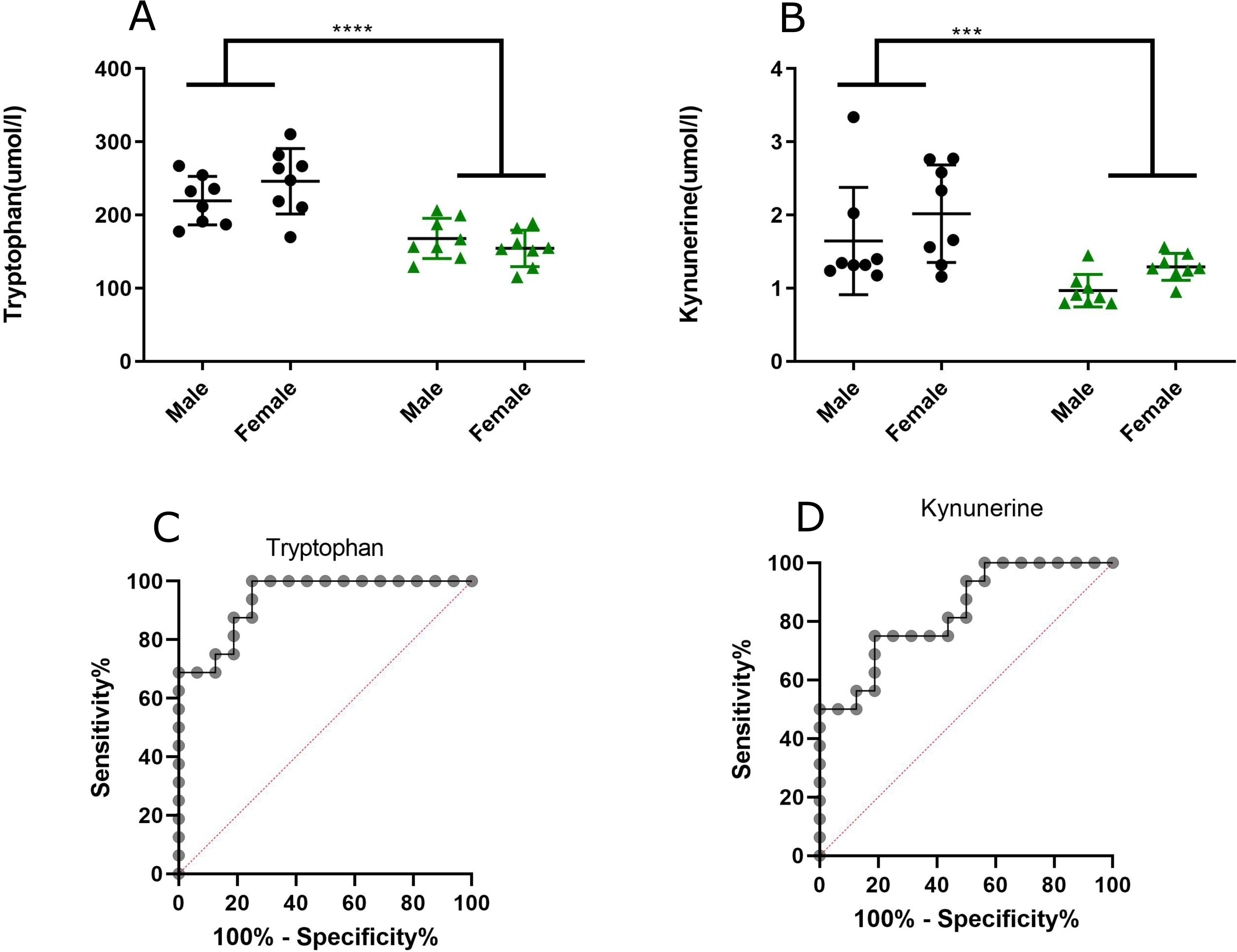
Targeted plasma metabolite profiling in mice at day 5 post-naproxen diet shows decrease in Tryptophan and Kynurenine. A) Plasma tryptophan B) Plasma Kynurenine C) ROC plot of tryptophan D) ROC plot of Kynurenine on day 5 post naproxen diet. The black and the green dots represent the control and the naproxen diet, respectively. The data is expressed as mean ± sd. * designates statistical significance, *** p≤ 0.001, **** p≤ 0.0001 respectively.

**fig. S5:**
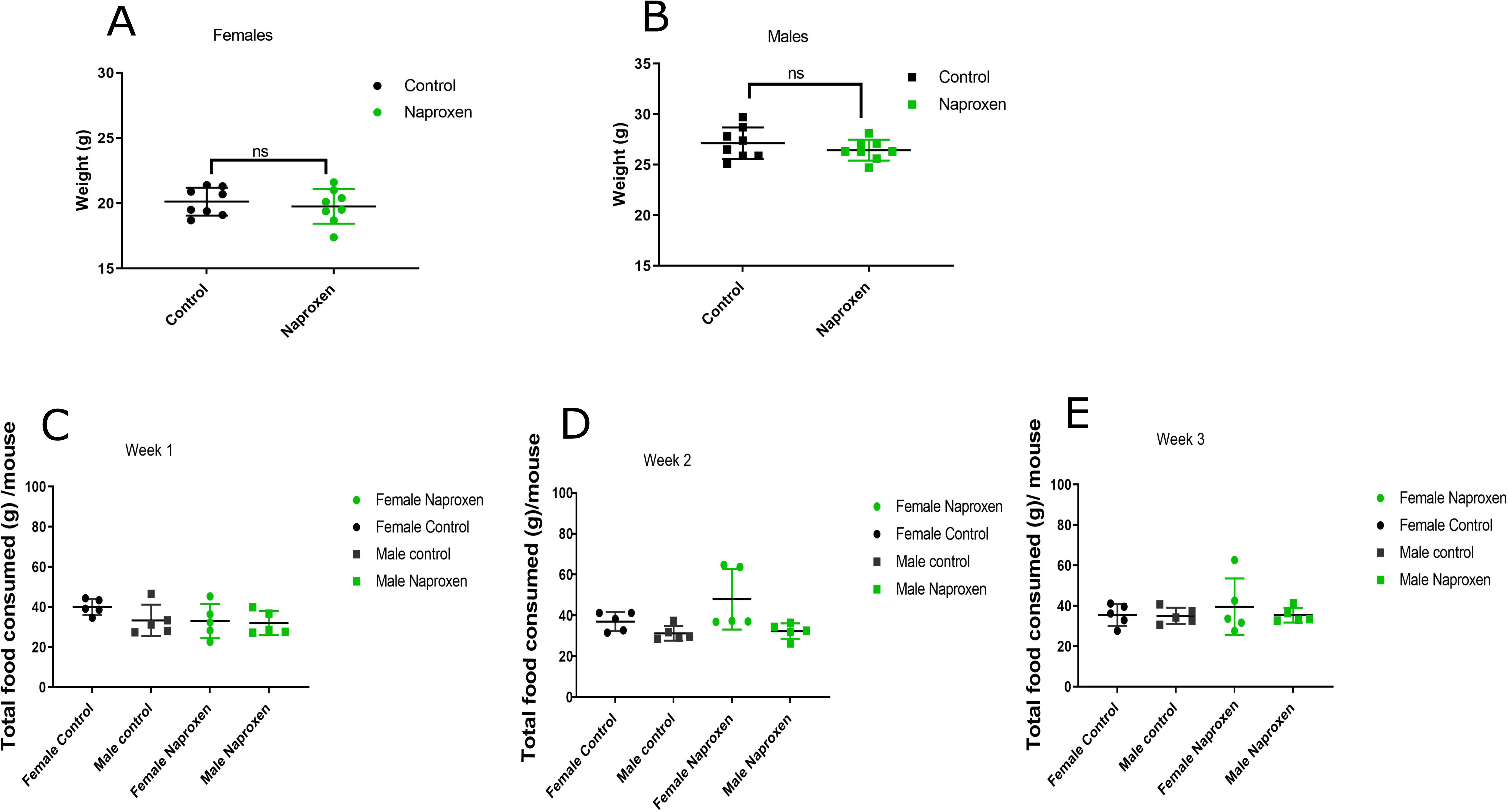
Weight and food intake in mice for three weeks in the Naproxen diet exhibit no difference between the groups. A) Weights of female mice in control chow and Naproxen diet after three weeks of diet. B) Weights of the male mice in the control chow and Naproxen diet after three weeks of diet. C) Food intake of the mice in week 1. D) Food intake of the mice in week 2. E) Food intake of the mice in week 3. The black and the green dots represent the control and Naproxen diet, respectively. The data is expressed as mean ± SD.

**fig. S6:**
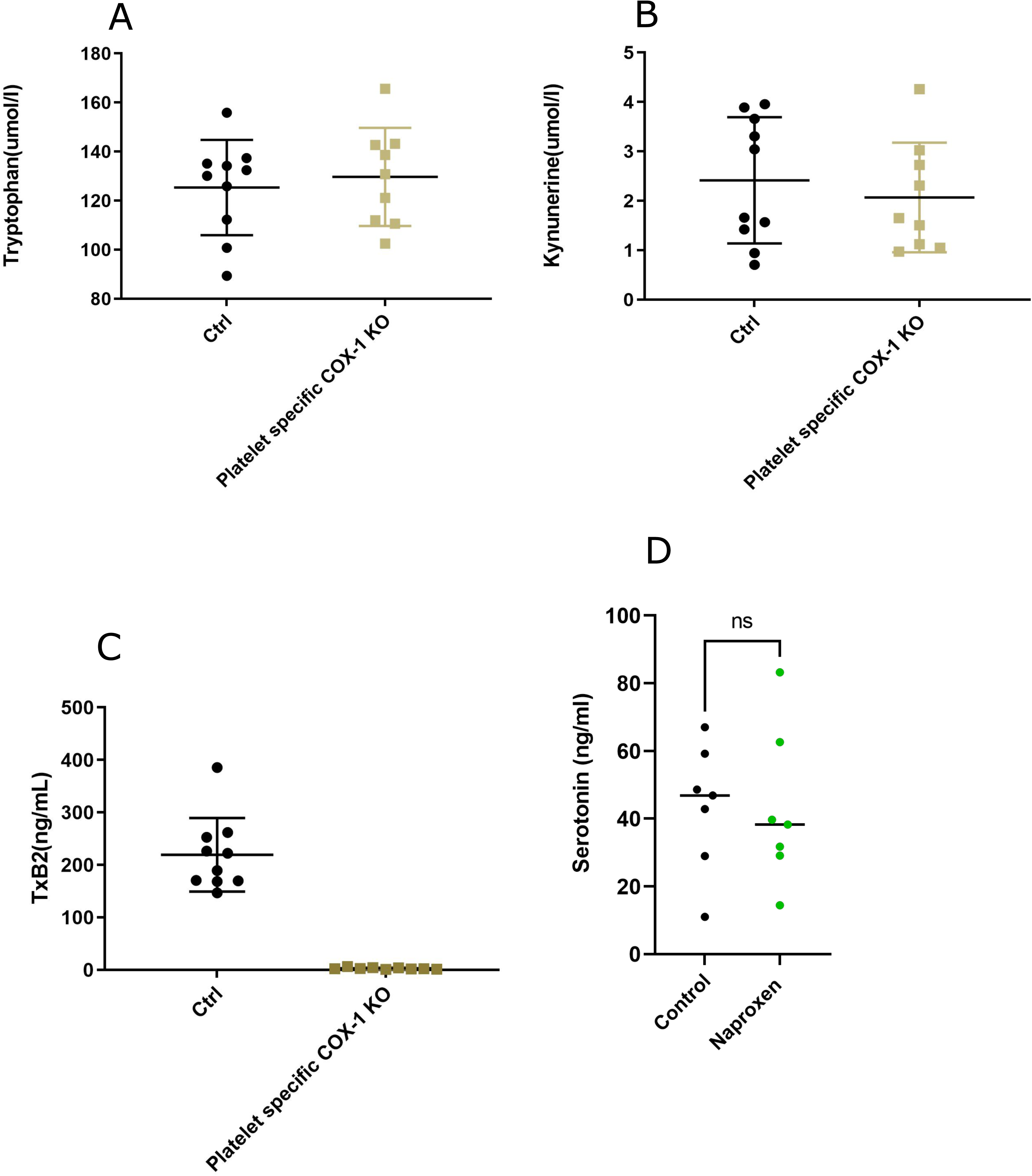
Targeted plasma metabolite profiling in Pf4-Δ Cre ^+/-^/COX-1fx/fx mice show no change with respect to control mice. A) Plasma Tryptophan B) Plasma Kynurenine C) Serum TxB2. The black dots and the yellow squares refer to controls and Pf4-Δ Cre ^+/-^/COX-1fx/fx respectively. The data is expressed as mean ± SD. D) Plasma Serotonin of C57Bl/6 in Control chow and Naproxen diet.

**fig. S7:**
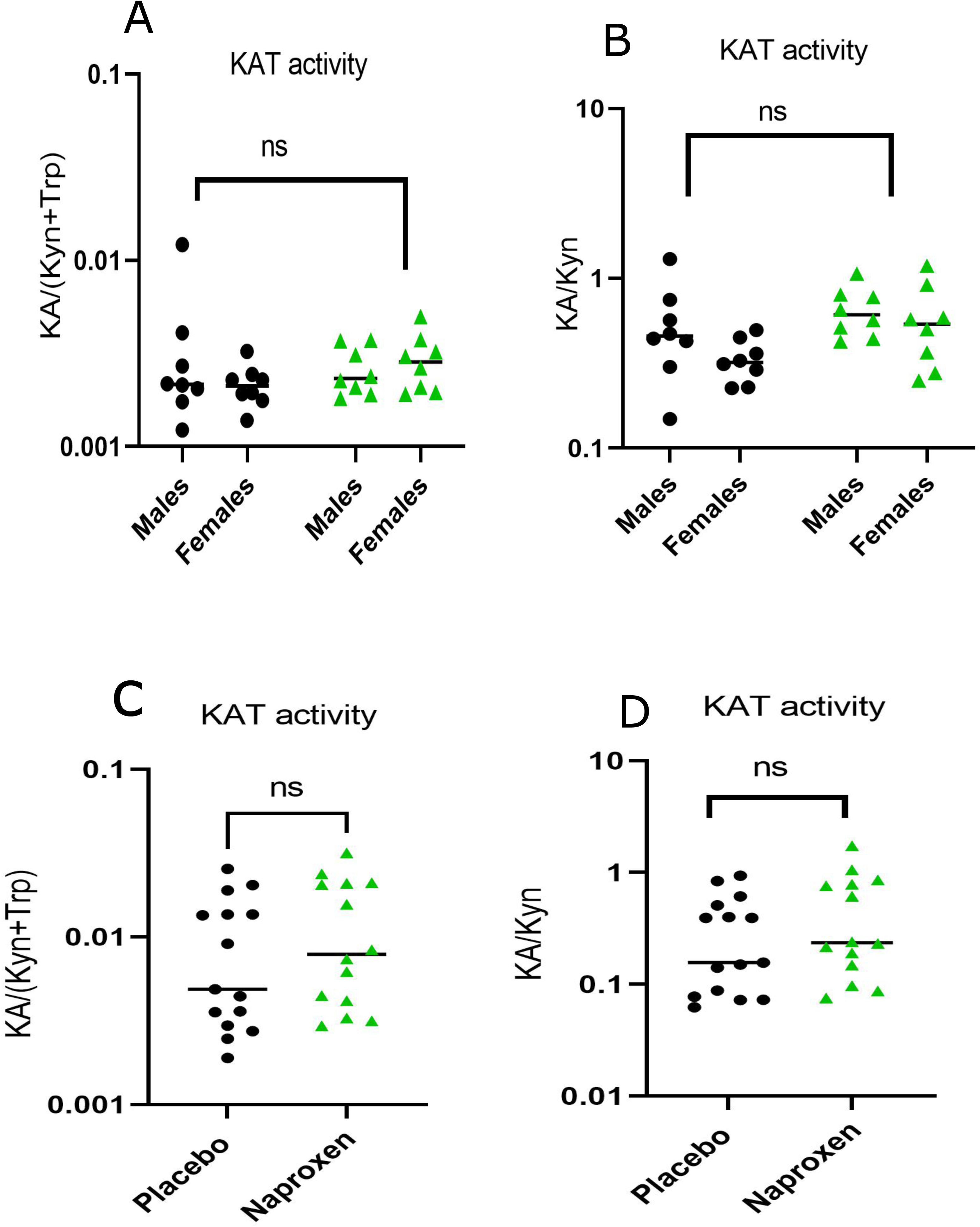
KAT activity is not altered by Naproxen treatment. A-B: KAT activity assessed in mice in control and Naproxen diet. C-D: KAT activity evaluated in human volunteers. Black dots and green triangles refer to control and Naproxen treatment, respectively.

**fig. S8:**
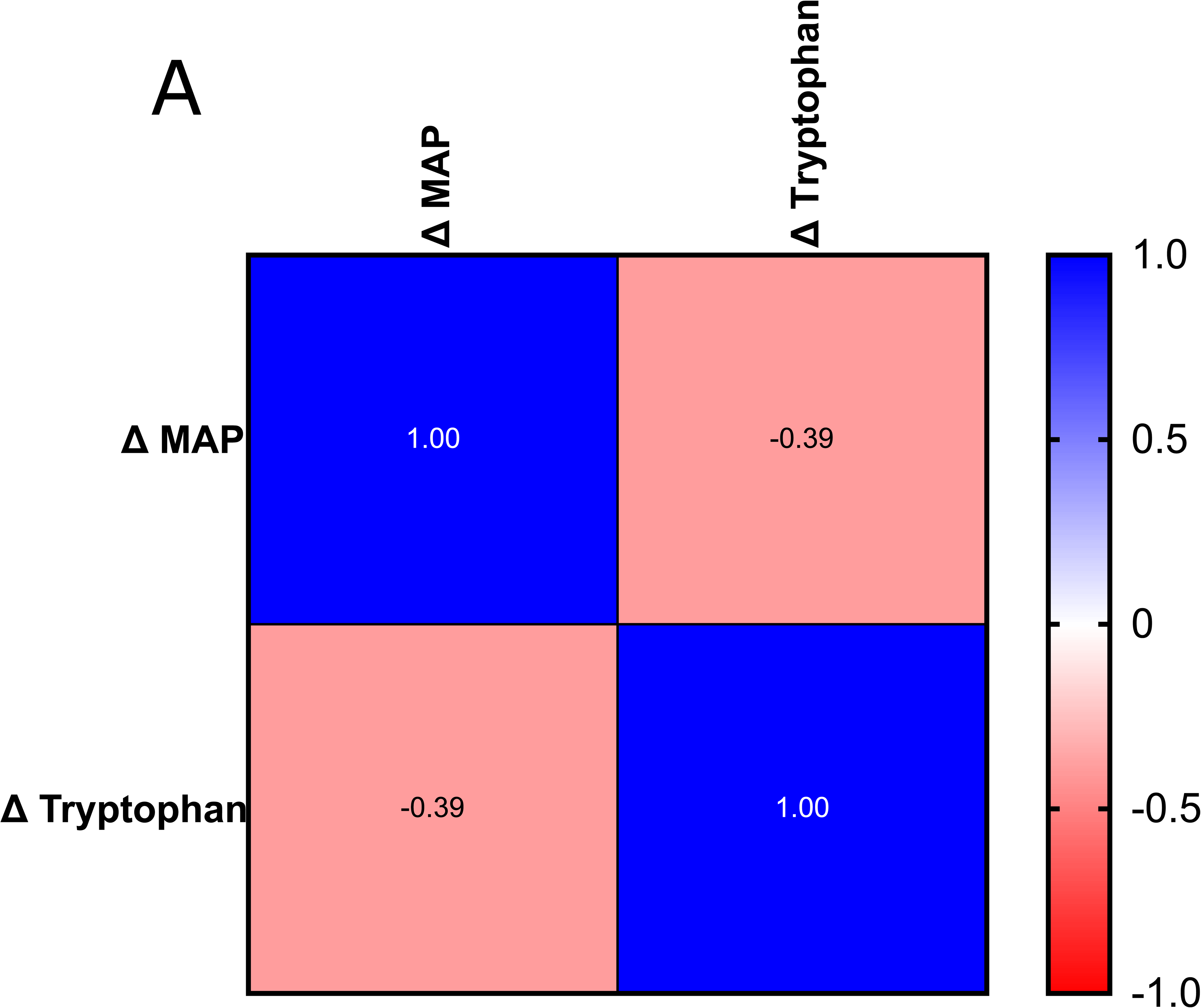
Tryptophan is related to cardiac-related phenotypes in Naproxen treatment. Spearman correlation between delta Mean Arterial Pressure (MAP) and delta plasma tryptophan levels in humans in Naproxen treatment.

**fig. S9:**
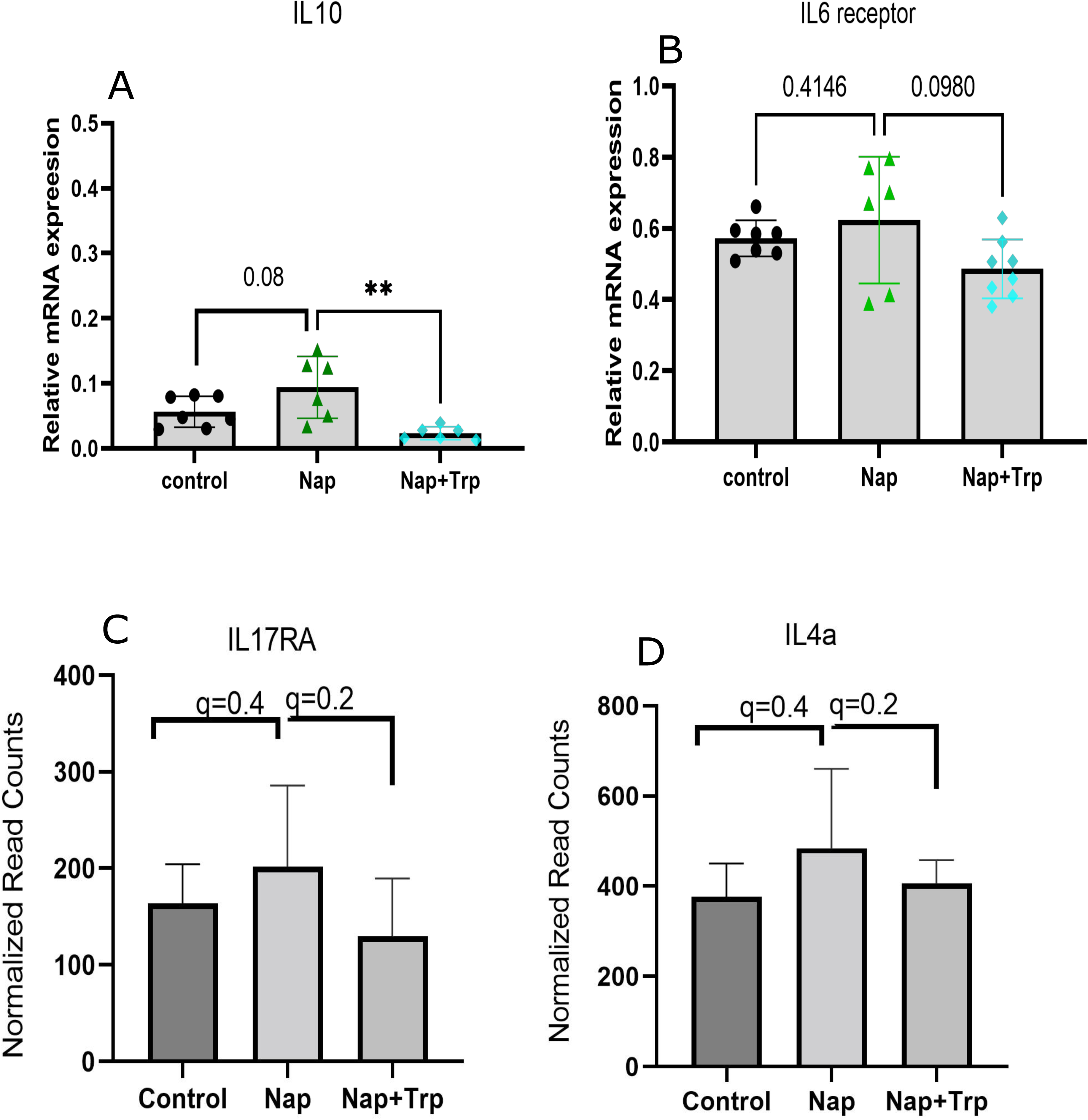
Tryptophan supplementation rescues cytokine profiles of heart in Naproxen treatment. A) IL-10 B) IL-6 Receptor C) IL-17RA D) IL-4a. The black circles, green triangles, and blue diamonds refer to control, naproxen and “naproxen+tryptophan” treatments. ** p≤ 0.01.

**fig. S10:**
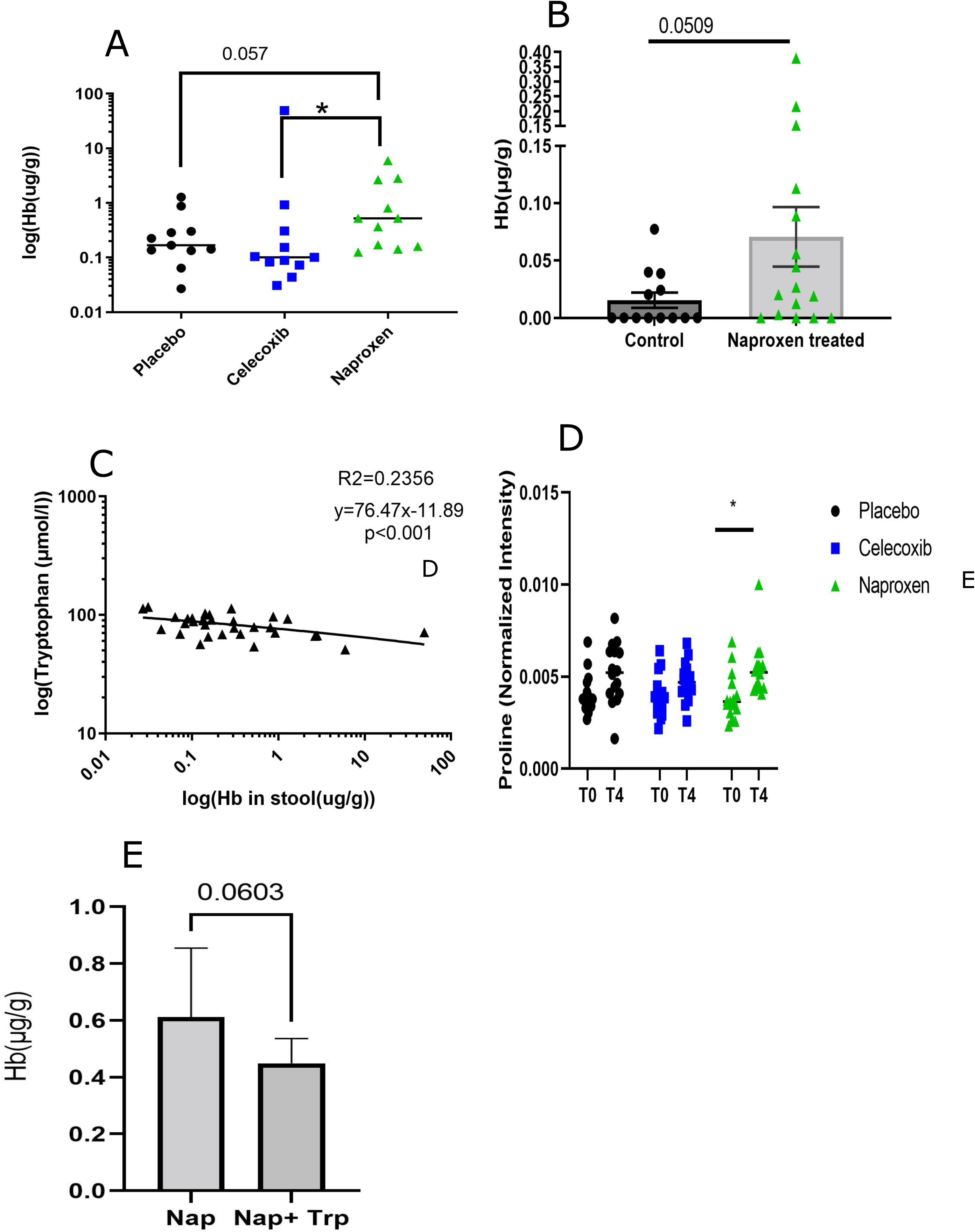
Tryptophan is related to Gut inflammation in Naproxen treatment. A) Hemoglobin content in Human stool in Placebo, Celecoxib, and Naproxen treatment. B) Hemoglobin content in mice stool measured in control and naproxen treatment of 3 weeks. C) Negative association of plasma tryptophan vs. Hemoglobin content in stool in humans. D) Plasma Proline concentration in Placebo, Celecoxib, and Naproxen. E) Hemoglobin content in mice stool after the tryptophan supplementation. The black dots, blue squares, and green triangles refer to placebo, celecoxib, and naproxen treatment. * designates statistical significance, * p≤ 0.05.

