## Supplementary Table 1 for "Metabolomic Response to Non-Steroidal Anti-Inflammatory Drugs"

Supplementary Table 1: Rescued Genes after Tryptophan Challenge

| **Gene_id** | **gene_name** | **gene_biotype** | **Trpnap_v_nap.q_value** |
| --- | --- | --- | --- |
| ENSMUSG00000022377 | Asap1 | protein_coding | 0.006818966 |
| ENSMUSG00000073414 | Mpig6b | protein_coding | 0.015367443 |
| ENSMUSG00000025804 | Ccr1 | protein_coding | 0.015588081 |
| ENSMUSG00000043740 | B430306N03Rik | protein_coding | 0.004284167 |
| ENSMUSG00000006642 | Tcf23 | protein_coding | 0.031731428 |
| ENSMUSG00000075224 | Lrrc55 | protein_coding | 0.054861429 |
| ENSMUSG00000025407 | Gli1 | protein_coding | 0.073937921 |
| ENSMUSG00000024222 | Fkbp5 | protein_coding | 0.024814073 |
| ENSMUSG00000075254 | Heg1 | protein_coding | 0.005422338 |
| ENSMUSG00000039031 | Arhgap18 | protein_coding | 0.004798355 |
| ENSMUSG00000020641 | Rsad2 | protein_coding | 0.021678581 |
| ENSMUSG00000066687 | Zbtb16 | protein_coding | 3.43E-09 |
| ENSMUSG00000022146 | Osmr | protein_coding | 0.03650584 |
| ENSMUSG00000025006 | Sorbs1 | protein_coding | 0.100270608 |
| ENSMUSG00000047592 | Nxpe5 | protein_coding | 0.017385166 |
| ENSMUSG00000047798 | Cd300lf | protein_coding | 0.107342425 |
| ENSMUSG00000024104 | Washc2 | protein_coding | 0.107575992 |
| ENSMUSG00000023216 | Epb42 | protein_coding | 0.109437589 |
| ENSMUSG00000030889 | Vwa3a | protein_coding | 0.114827813 |
| ENSMUSG00000018604 | Tbx3 | protein_coding | 0.120466524 |
| ENSMUSG00000020227 | Irak3 | protein_coding | 0.037974605 |
| ENSMUSG00000058624 | Gda | protein_coding | 0.081017768 |
| ENSMUSG00000043079 | Synpo | protein_coding | 0.124514428 |
| ENSMUSG00000025584 | Pde8a | protein_coding | 0.045463597 |
| ENSMUSG00000052749 | Trim30b | protein_coding | 0.135817432 |
| ENSMUSG00000091694 | Apol11b | protein_coding | 0.141484984 |
| ENSMUSG00000026890 | Lhx6 | protein_coding | 0.043458087 |
| ENSMUSG00000026971 | Itgb6 | protein_coding | 0.00299495 |
| ENSMUSG00000066877 | Nck2 | protein_coding | 0.096117263 |
| ENSMUSG00000031762 | Mt2 | protein_coding | 0.143565557 |
| ENSMUSG00000021591 | Glrx | protein_coding | 0.145152904 |
| ENSMUSG00000027630 | Tbl1xr1 | protein_coding | 0.083472922 |
| ENSMUSG00000029417 | Cxcl9 | protein_coding | 0.155919628 |
| ENSMUSG00000028906 | Epb41 | protein_coding | 0.06461216 |
| ENSMUSG00000031302 | Nlgn3 | protein_coding | 0.168832338 |
| ENSMUSG00000041684 | Bivm | protein_coding | 0.006845572 |
| ENSMUSG00000051839 | Gypa | protein_coding | 0.170371858 |
| ENSMUSG00000073400 | Trim10 | protein_coding | 0.170763587 |
| ENSMUSG00000017861 | Mybl2 | protein_coding | 0.172912239 |
| ENSMUSG00000024679 | Ms4a6d | protein_coding | 0.155919628 |
| ENSMUSG00000024935 | Slc1a1 | protein_coding | 0.180569777 |
| ENSMUSG00000022220 | Adcy4 | protein_coding | 0.043458087 |
| ENSMUSG00000045201 | Lrrc3b | protein_coding | 0.188917581 |
| ENSMUSG00000073879 | Gm5859 | unprocessed_pseudogene | 0.188917581 |
| ENSMUSG00000034135 | Sik3 | protein_coding | 0.189926197 |
| ENSMUSG00000024481 | Lvrn | protein_coding | 0.176430058 |
| ENSMUSG00000030655 | Smg1 | protein_coding | 0.10704898 |
| ENSMUSG00000051650 | B3gnt2 | protein_coding | 0.191305244 |
| ENSMUSG00000034684 | Sema3f | protein_coding | 0.155919628 |
| ENSMUSG00000041920 | Slc16a6 | protein_coding | 0.195630299 |
| ENSMUSG00000046562 | Unc119b | protein_coding | 0.195647971 |
| ENSMUSG00000015957 | Wnt11 | protein_coding | 0.195961685 |
| ENSMUSG00000034157 | Cipc | protein_coding | 0.201487752 |
| ENSMUSG00000025013 | Tll2 | protein_coding | 0.007022733 |
| ENSMUSG00000095794 | Igkv6-17 | IG_V_gene | 0.132471092 |
| ENSMUSG00000005667 | Mthfd2 | protein_coding | 0.204265439 |
| ENSMUSG00000032508 | Myd88 | protein_coding | 0.205619158 |
| ENSMUSG00000009185 | Ccl8 | protein_coding | 0.054916888 |
| ENSMUSG00000045328 | Cenpe | protein_coding | 0.211998298 |
| ENSMUSG00000020185 | E2f7 | protein_coding | 0.213277812 |
| ENSMUSG00000003526 | Prodh | protein_coding | 0.213846215 |
| ENSMUSG00000035835 | Plppr3 | protein_coding | 0.0883098 |
| ENSMUSG00000045555 | Mettl24 | protein_coding | 0.195630299 |
| ENSMUSG00000031766 | Slc12a3 | protein_coding | 0.01090889 |
| ENSMUSG00000111389 | Gm39465 | lincRNA | 0.059124183 |
| ENSMUSG00000042331 | Specc1 | protein_coding | 0.216148223 |
| ENSMUSG00000051456 | Hspb3 | protein_coding | 0.221021676 |
| ENSMUSG00000060548 | Tnfrsf19 | protein_coding | 0.225276837 |
| ENSMUSG00000044456 | Rin3 | protein_coding | 0.228469462 |
| ENSMUSG00000053297 | AI854703 | protein_coding | 0.093242904 |
| ENSMUSG00000047953 | Gp5 | protein_coding | 0.11060791 |
| ENSMUSG00000053687 | Dpep2 | protein_coding | 0.242278103 |
| ENSMUSG00000030748 | Il4ra | protein_coding | 0.243030226 |
| ENSMUSG00000045954 | Cavin2 | protein_coding | 0.116577745 |
| ENSMUSG00000054203 | Ifi205 | protein_coding | 0.192533693 |
| ENSMUSG00000024081 | Cebpz | protein_coding | 0.247056129 |
| ENSMUSG00000043259 | Fam13c | protein_coding | 0.247056129 |
| ENSMUSG00000006574 | Slc4a1 | protein_coding | 0.248618962 |
| ENSMUSG00000024043 | Arhgap28 | protein_coding | 0.253880919 |
| ENSMUSG00000005413 | Hmox1 | protein_coding | 0.201828545 |
| ENSMUSG00000040485 | Lrrc52 | protein_coding | 0.260180445 |
| ENSMUSG00000020262 | Adarb1 | protein_coding | 0.017890937 |
| ENSMUSG00000022469 | Rapgef3 | protein_coding | 0.22932802 |
| ENSMUSG00000024085 | Man2a1 | protein_coding | 0.010908263 |
| ENSMUSG00000031990 | Jam3 | protein_coding | 0.122607426 |
| ENSMUSG00000078897 | Gm4724 | protein_coding | 0.189528793 |
| ENSMUSG00000004709 | Cd244a | protein_coding | 0.251541643 |
| ENSMUSG00000034616 | Ssh3 | protein_coding | 0.269839143 |
| ENSMUSG00000027463 | Slc52a3 | protein_coding | 0.272904493 |
| ENSMUSG00000033024 | Klra9 | protein_coding | 0.272904493 |
| ENSMUSG00000044468 | Tent5c | protein_coding | 0.272904493 |
| ENSMUSG00000047205 | Dusp18 | protein_coding | 0.273988411 |
| ENSMUSG00000004552 | Ctse | protein_coding | 0.279121927 |
| ENSMUSG00000035849 | Krt222 | protein_coding | 0.195630299 |
| ENSMUSG00000081822 | Gm15626 | processed_pseudogene | 0.004462429 |
| ENSMUSG00000000732 | Icosl | protein_coding | 0.261572224 |
| ENSMUSG00000026288 | Inpp5d | protein_coding | 0.195630299 |
| ENSMUSG00000033174 | Mgll | protein_coding | 0.286282957 |
| ENSMUSG00000026536 | Ifi211 | protein_coding | 0.288174397 |
| ENSMUSG00000086040 | Wipf3 | protein_coding | 0.144851047 |
| ENSMUSG00000049723 | Mmp12 | protein_coding | 0.060938719 |
| ENSMUSG00000026180 | Cxcr2 | protein_coding | 0.291956965 |
| ENSMUSG00000096488 | Gm10409 | protein_coding | 0.291956965 |
| ENSMUSG00000046201 | Scaf8 | protein_coding | 0.073937921 |
| ENSMUSG00000013846 | St3gal1 | protein_coding | 0.017120226 |
| ENSMUSG00000002147 | Stat6 | protein_coding | 0.296785396 |
| ENSMUSG00000031154 | Otud5 | protein_coding | 0.217333262 |
| ENSMUSG00000028776 | Tinagl1 | protein_coding | 0.301637264 |
| ENSMUSG00000025092 | Hspa12a | protein_coding | 0.195961685 |
| ENSMUSG00000035992 | Fnip1 | protein_coding | 0.11922112 |
| ENSMUSG00000075602 | Ly6a | protein_coding | 0.258102791 |
| ENSMUSG00000063060 | Sox7 | protein_coding | 0.307327613 |
| ENSMUSG00000030283 | St8sia1 | protein_coding | 0.308043084 |
| ENSMUSG00000023993 | Treml1 | protein_coding | 0.278726146 |
| ENSMUSG00000050592 | Fam78a | protein_coding | 0.181575406 |
| ENSMUSG00000022999 | Lmbr1l | protein_coding | 0.308797588 |
| ENSMUSG00000078763 | Slfn1 | protein_coding | 0.308797588 |
| ENSMUSG00000030852 | Tacc2 | protein_coding | 0.069449209 |
| ENSMUSG00000071714 | Csf2rb2 | protein_coding | 0.155919628 |
| ENSMUSG00000026579 | F5 | protein_coding | 0.194516757 |
| ENSMUSG00000072571 | Tmem253 | protein_coding | 0.217333262 |
| ENSMUSG00000036611 | Eepd1 | protein_coding | 0.238753426 |
| ENSMUSG00000037965 | Zc3h7a | protein_coding | 0.038834144 |
| ENSMUSG00000049588 | Ccdc69 | protein_coding | 0.316715986 |
| ENSMUSG00000026779 | Mastl | protein_coding | 0.11405541 |
| ENSMUSG00000031776 | Arl2bp | protein_coding | 0.211250803 |
| ENSMUSG00000035863 | Palm | protein_coding | 0.013856846 |
| ENSMUSG00000048582 | Gja3 | protein_coding | 0.210166741 |
| ENSMUSG00000050675 | Gp1ba | protein_coding | 0.173189155 |
| ENSMUSG00000055639 | Dach1 | protein_coding | 0.170164084 |
| ENSMUSG00000097649 | Gm10561 | antisense | 0.203482466 |
| ENSMUSG00000025597 | Klhl4 | protein_coding | 0.157072083 |
| ENSMUSG00000110631 | Gm42047 | lincRNA | 0.242653287 |
| ENSMUSG00000025239 | Limd1 | protein_coding | 0.306748088 |
| ENSMUSG00000058794 | Nfe2 | protein_coding | 0.124514428 |
| ENSMUSG00000027955 | Gask1b | protein_coding | 0.323808365 |
| ENSMUSG00000032418 | Me1 | protein_coding | 0.289477643 |
| ENSMUSG00000090394 | 4930523C07Rik | protein_coding | 0.331593667 |
| ENSMUSG00000027293 | Ehd4 | protein_coding | 0.167590673 |
| ENSMUSG00000046470 | Sox18 | protein_coding | 0.335075054 |
| ENSMUSG00000018983 | E2f2 | protein_coding | 0.339260809 |
| ENSMUSG00000020034 | Tcp11l2 | protein_coding | 0.023849396 |
| ENSMUSG00000024471 | Myot | protein_coding | 0.053387454 |
| ENSMUSG00000024044 | Epb41l3 | protein_coding | 0.265281028 |
| ENSMUSG00000032400 | Zwilch | protein_coding | 0.273988411 |
| ENSMUSG00000034903 | Cobll1 | protein_coding | 0.102877902 |
| ENSMUSG00000024558 | Mapk4 | protein_coding | 0.317552839 |
| ENSMUSG00000020689 | Itgb3 | protein_coding | 0.054916888 |
| ENSMUSG00000020787 | P2rx1 | protein_coding | 0.341057038 |
| ENSMUSG00000022770 | Dlg1 | protein_coding | 0.057107458 |
| ENSMUSG00000028874 | Fgr | protein_coding | 0.216148223 |
| ENSMUSG00000033159 | Cnppd1 | protein_coding | 0.176430058 |
| ENSMUSG00000056214 | Pard6g | protein_coding | 0.291986697 |
| ENSMUSG00000116506 | 5730414N17Rik | lincRNA | 0.206377569 |
| ENSMUSG00000074743 | Thbd | protein_coding | 0.086208066 |
| ENSMUSG00000040536 | Necab1 | protein_coding | 0.346410891 |
| ENSMUSG00000047878 | A4galt | protein_coding | 0.20902158 |
| ENSMUSG00000038007 | Acer2 | protein_coding | 0.346742386 |
| ENSMUSG00000028859 | Csf3r | protein_coding | 0.352598749 |
| ENSMUSG00000000204 | Slfn4 | protein_coding | 0.353087064 |
| ENSMUSG00000038264 | Sema7a | protein_coding | 0.296819241 |
| ENSMUSG00000004698 | Hdac9 | protein_coding | 0.02241063 |
| ENSMUSG00000026656 | Fcgr2b | protein_coding | 0.195961685 |
| ENSMUSG00000026070 | Il18r1 | protein_coding | 0.333499974 |
| ENSMUSG00000040441 | Slc26a10 | protein_coding | 0.362803729 |
| ENSMUSG00000049303 | Syt12 | protein_coding | 0.034674361 |
| ENSMUSG00000074677 | Sirpb1c | protein_coding | 0.155919628 |
| ENSMUSG00000112707 | D830005E20Rik | antisense | 0.367066085 |
| ENSMUSG00000034898 | Filip1 | protein_coding | 0.010375623 |
| ENSMUSG00000026640 | Plxna2 | protein_coding | 0.093242904 |
| ENSMUSG00000031803 | B3gnt3 | protein_coding | 0.261572224 |
| ENSMUSG00000098708 | Gm27252 | processed_transcript | 0.334365224 |
| ENSMUSG00000033170 | Card10 | protein_coding | 0.015588081 |
| ENSMUSG00000009292 | Trpm2 | protein_coding | 0.373011944 |
| ENSMUSG00000048120 | Entpd1 | protein_coding | 0.318857863 |
| ENSMUSG00000019779 | Frk | protein_coding | 0.374795015 |
| ENSMUSG00000022623 | Shank3 | protein_coding | 0.373277016 |
| ENSMUSG00000020044 | Timp3 | protein_coding | 0.195630299 |
| ENSMUSG00000029777 | Gars | protein_coding | 0.337211797 |
| ENSMUSG00000079164 | Tlr5 | protein_coding | 0.375820672 |
| ENSMUSG00000019866 | Crybg1 | protein_coding | 0.0883098 |
| ENSMUSG00000020914 | Top2a | protein_coding | 0.377312174 |
| ENSMUSG00000031986 | Sprtn | protein_coding | 0.380136827 |
| ENSMUSG00000053113 | Socs3 | protein_coding | 0.16480345 |
| ENSMUSG00000042742 | Bmt2 | protein_coding | 0.383095527 |
| ENSMUSG00000037820 | Tgm2 | protein_coding | 0.385209013 |
| ENSMUSG00000071713 | Csf2rb | protein_coding | 0.210447565 |
| ENSMUSG00000020160 | Meis1 | protein_coding | 0.38595696 |
| ENSMUSG00000022974 | Paxbp1 | protein_coding | 0.11060791 |
| ENSMUSG00000002797 | Ggct | protein_coding | 0.234140562 |
| ENSMUSG00000009687 | Fxyd5 | protein_coding | 0.155919628 |
| ENSMUSG00000028184 | Adgrl2 | protein_coding | 0.37275084 |
| ENSMUSG00000042428 | Mgat3 | protein_coding | 0.143565557 |
| ENSMUSG00000051169 | Rpusd3 | protein_coding | 0.195961685 |
| ENSMUSG00000059436 | Max | protein_coding | 0.386598412 |
| ENSMUSG00000042182 | Bend6 | protein_coding | 0.387400534 |
| ENSMUSG00000071454 | Dtnb | protein_coding | 0.388280466 |
| ENSMUSG00000033595 | Lgi3 | protein_coding | 0.388330583 |
| ENSMUSG00000034648 | Lrrn1 | protein_coding | 0.388478213 |
| ENSMUSG00000034858 | Fam214a | protein_coding | 0.073937921 |
| ENSMUSG00000020330 | Hmmr | protein_coding | 0.390634422 |
| ENSMUSG00000046818 | Ddit4l | protein_coding | 0.390766292 |
| ENSMUSG00000030054 | Gp9 | protein_coding | 0.392865608 |
| ENSMUSG00000025983 | Ccdc150 | protein_coding | 0.261572224 |
| ENSMUSG00000056069 | Otulinl | protein_coding | 0.060445942 |
| ENSMUSG00000027469 | Tpx2 | protein_coding | 0.383781904 |
| ENSMUSG00000042807 | Hecw2 | protein_coding | 0.155764341 |
| ENSMUSG00000048756 | Foxo3 | protein_coding | 0.191812893 |
| ENSMUSG00000072623 | Zfp9 | protein_coding | 0.399314024 |
| ENSMUSG00000021260 | Hhipl1 | protein_coding | 0.399367433 |
| ENSMUSG00000028862 | Map3k6 | protein_coding | 0.399659478 |
| ENSMUSG00000003934 | Efnb3 | protein_coding | 0.301496678 |
